# Low-dimensional encoding of decisions in parietal cortex reflects long-term training history

**DOI:** 10.1101/2021.10.07.463576

**Authors:** Kenneth W. Latimer, David J. Freedman

**Affiliations:** Department of Neurobiology, University of Chicago

## Abstract

Neurons in parietal cortex exhibit task-related activity during decision-making tasks. However, it remains unclear how long-term training to perform different tasks over months or even years shapes neural computations and representations. We examine lateral intraparietal area (LIP) responses during a visual motion delayed-match-to-category (DMC) task. We consider two pairs of monkeys with different training histories: one trained only on the DMC task, and another first trained to perform fine motion-direction discrimination. We introduce generalized multilinear models to quantify low-dimensional, task-relevant components in population activity. During the DMC task, we found stronger cosine-like motion-direction tuning in the pretrained monkeys than in the DMC-only monkeys, and that the pretrained monkeys’ performance depended more heavily on sample-test stimulus similarity. These results suggest that sensory representations in LIP depend on the sequence of tasks that the animals have learned, underscoring the importance of training history in studies with complex behavioral tasks.

## 1 Introduction

Activity of single neurons in the macaque lateral intraparietal area (LIP) encodes task-relevant information in a variety of decision-making tasks (Freedman & Ibos, 2018). As a result, LIP has been proposed to support many different neural computations underlying perceptual decision making, including abstract visual categorization (Freedman & Assad, 2016; Huk et al., 2017). Throughout a lifetime, animals learn to make many different kinds of decisions in a variety of tasks and contexts, and different animals collect a unique set of experiences that shape their perceptual and decision-making skills and strategies (Summerfield & De Lange, 2014; Goldstone & Byrge, 2015). In contrast, experiments designed to study neural mechanisms of decision making often focus on neurons recorded during a specific task in isolation. However, previously learned neural representations and strategies may impact how a cortical region is recruited when learning a new task. To understand the generality and flexibility of neural representations which support decision making, we aim to compare decision-related LIP activity in animals performing the same tasks, but with different long-term training histories.

We examine LIP recordings in four monkeys performing two related tasks in which they were required to determine if sequentially presented motion directions matched according to a learned rule (Fig. 1A). In both tasks, the monkey views two random dot motion stimuli (sample and test) separated by a delay period. To receive a reward, the animal responds by releasing a touch bar if the two stimuli match or by continuing to hold the touch bar on non-match trials. The delayed-match-to-sample (DMS) task is a memory-based, fine-direction discrimination task in which the sample and test motion stimuli match only if they are in the exact same direction. In the delayed-match-to-category (DMC) task, the stimuli match if the directions belong to the same category (red or blue) according to a learned arbitrary category rule. In the DMC task, two matching stimuli may be nearly 180° apart but belong to matching categories, while neighboring directions on different sides of the category boundary do not match. Thus, while the tasks use the same structure and stimuli, they require performing different perceptual and/or cognitive computations.

**Figure 1:**
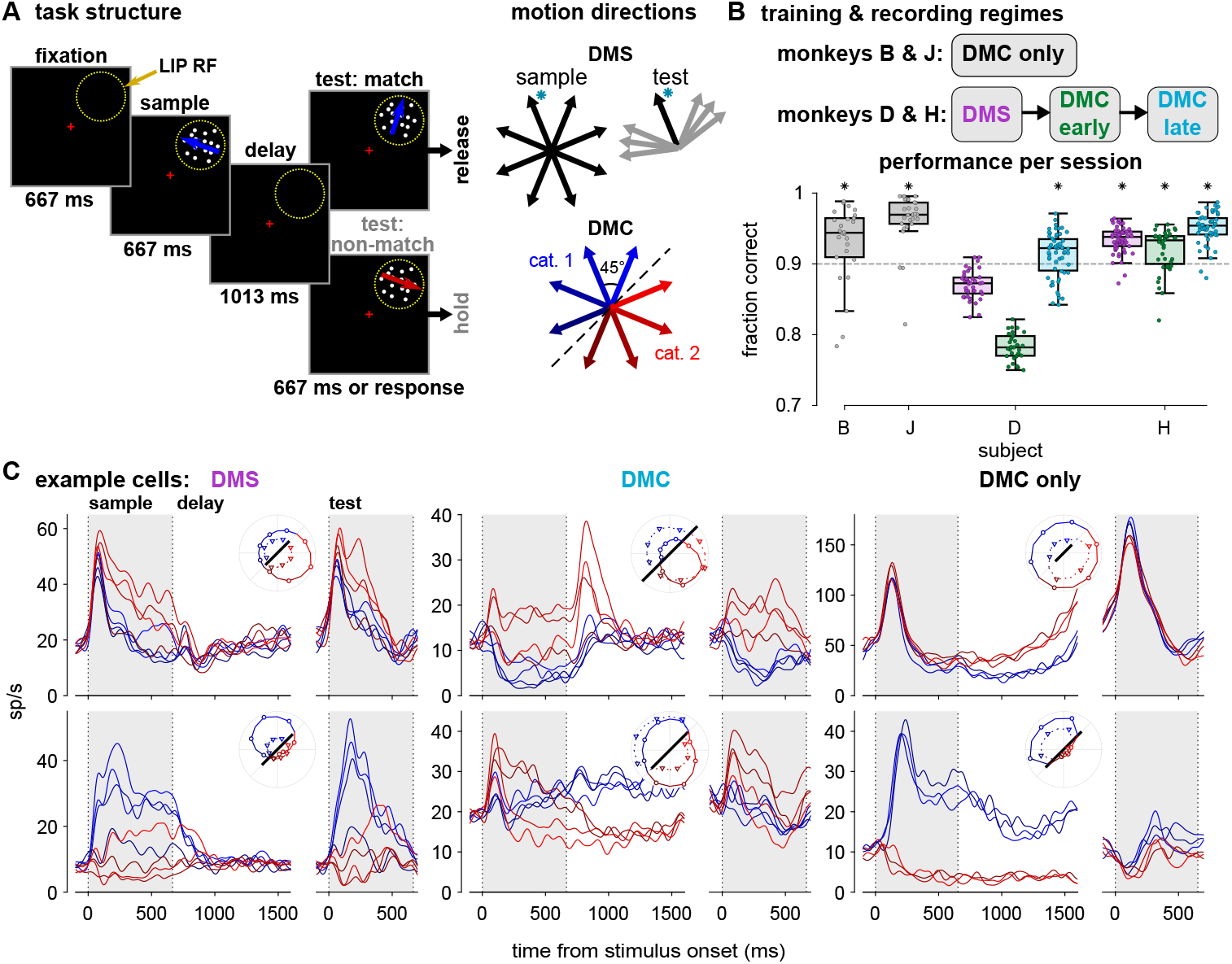
LIP recordings during DMS and DMC tasks. (**A**) In both tasks, the animal fixated and viewed a motion direction stimulus (sample). Following a delay period, a second stimulus (test) was presented. The monkey signaled if the sample and test stimuli matched by releasing a touch bar, otherwise the monkey was required to hold the touch bar. In the DMS task, the sample and test stimuli matched only if the directions were exactly the same. In the DMC task, the stimuli matched if they belonged to the same category: the motion directions were split into two equally sized categories with a 45°-225° boundary (red and blue directions; the boundary was constant for all sessions). The motion stimuli were placed inside the LIP cell’s response field (yellow circle) during recording. (**B**) (top) Training and recording regimes for the four monkeys. (bottom) Performance during each recording session (dots) for each animal are summarized by the box plots. Colors correspond to the task and training period (DMS, DMC early or late, and DMC only). Asterisks indicate median per-session performance is greater than 90 % (*p* < 0.01, one-sided sign test, Holm-Bonferroni corrected). All four monkeys learned to perform the DMC task with a median performance of at least 90 % per session. (**C**) Mean firing rates of six single LIP cells recorded during each task. Colors correspond to the stimulus direction and category. Firing rates aligned to the sample stimulus onset are averaged by sample direction (left), and the test stimulus aligned rates are averaged over test direction (right). Although motion categories were not part of the DMS task, the directions are labeled blue or red for consistency. (inset polar plots) The mean firing rate for each sample direction during the sample stimulus presentation (circles and solid lines; 0 ms to 650 ms after motion onset) and delay period (triangles and dotted lines; 800 ms to 1450 ms after motion onset). The solid black line denotes 20 sp/s.

We consider two pairs of monkeys with two different training histories (Fig. 1B). In one pair of monkeys (B and J; Swaminathan & Freedman, 2012), the monkeys were trained only to perform the DMC task (i.e., without first training the monkeys on fine discrimination), and LIP recordings were made after training was completed (DMC only populations). The second pair of monkeys (pretrained monkeys D and H; Sarma et al., 2016), was first trained extensively on the DMS task, and LIP recordings were obtained after training (DMS population). The monkeys were then retrained on the DMC task, and a set of LIP recordings was made during an intermediate stage of training (when the monkeys’ performance stabilized; DMC early populations). After the DMC-early recordings, the monkeys received additional training which overemphasized near-category-boundary sample stimuli (the most difficult conditions where the monkeys’ performance was lowest) so that the monkeys’ performance increased. After the second training stage was complete, a final set of LIP neurons was recorded during the DMC task (DMC late populations). In this study, the monkeys did not perform both tasks during a single session; they were switched exclusively to the DMC task and retrained over the course of months. In total, we analyzed eight LIP populations from four animals.

Not only do the DMS and DMC tasks share the same structure, timings, and stimuli, many of the sample-test pairs are rewarded for the same responses in both contexts (e.g., sample and test stimuli of the same direction match in both tasks). It is plausible that pretrained monkeys may reuse strategies acquired for performing the discrimination task while learning the DMC task. Similarly, different training histories may lead to different behavioral strategies to perform the DMC task, and different strategies may give rise to different patterns of activity in LIP (Tang et al., 2020). Previous studies have found that stimulus encoding and working-memory dependent sustained-firing activity in the prefrontal cortex during cognitive tasks depends on training (Qi & Constantinidis, 2013; Li et al., 2020). Additionally, the causal contribution of the middle temporal (MT) area of macaque visual cortex, which encodes motion and projects to LIP (Born & Bradley, 2005), depends on training history during motion-direction discrimination (Liu & Pack, 2017) and coarse depth discrimination (Chowdhury & DeAngelis, 2008). We therefore hypothesize that differences in training history could result in differences in LIP population activity during the DMC task which reflect behaviorally relevant aspects of the neural computations underlying categorization (Churchland & Kiani, 2016).

Direction and category selectivity is visible in the average firing rates of single LIP neurons in both pairs of monkeys during the DMC task (Fig. 1C). However, based on single cells alone it is difficult to uncover the computations involved in the DMC task: how sample category is computed and then stored during the delay period, or how the test stimulus is compared to the sample. Instead, we take a dimensionality reduction approach to compare the low-dimensional geometry of population responses to better illuminate how LIP encodes different tasks (Okazawa et al., 2021).

While many methods of dimensionality reduction are available, we sought a compact, low-dimensional description of LIP responses that quantified the population responses as a function of the task variables. Moreover, we wished to perform dimensionality reduction on the trial-by-trial spike train responses (as opposed to trial-averaged spike rates) within each population in order to account for structure in the neural activity beyond mean firing rates (e.g., bursting or oscillations). We therefore introduce the generalized multilinear model (GMLM) as a model-based dimensionality reduction method for population activity during flexible cognitive tasks (Fig. 2A). The GMLM is a tensor-regression extension of the generalized linear model (GLM) which describes a single neuron’s spiking response to different task events through a set of linear weights or kernels (Zhou et al., 2013; Park et al., 2014; Robinson et al., 2016; Kossaifi et al., 2020). The GMLM fits the data from all cells in a dataset into one compact representation by taking a low-rank tensor decomposition of linear kernels of the task events — the sample and test stimuli and the touch-bar release — that best describes the shared response dynamics across the entire population as a function of the task variables. This contrasts with the GLM, which fits each cell individually without directly recovering low-dimensional structure. Similarly to exponential family principal components analysis (Mohamed et al., 2008), the GMLM can be applied directly to binned spike count data rather than smoothed spike rates. The GMLM inherits the GLM’s flexibility for modeling trials with variable structure. For example, the timing of the end of the trial is controlled by the animal via their releasing the touch-bar. In contrast, dimensionality reduction approaches based on peristimulus time histograms (PSTH) require temporally aligned trials (e.g., principal components analysis-based methods; Kobak et al., 2016; Aoi et al., 2020), thereby limiting those approaches’ ability to quantify motion and category tuning or touch-bar response related activity during the test period. Stimulus category and direction are low-dimensional variables, and motion direction tuning in sensory regions such as area MT can be well-captured by simple parametric models (Rust et al., 2006). Therefore, the GMLM is well-suited for modeling how LIP populations represent combinations of these task variables during the DMC task.

**Figure 2:**
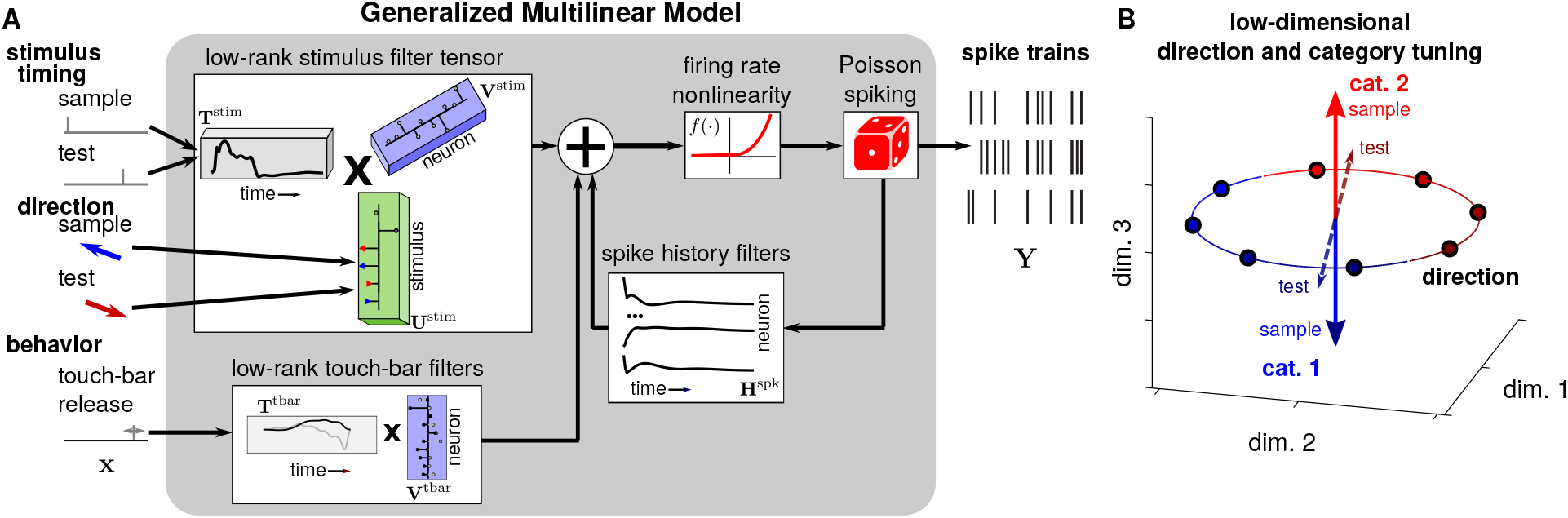
The generalized multilinear model for dimensionality reduction of neural populations. (**A**) Diagram of the GMLM. Incoming stimuli are factorized into temporal events and stimulus weights that encode direction and category information (Fig. S1E-I). A set of temporal kernels and stimulus coefficients filter the stimuli into a low-dimensional stimulus response space. The touch-bar release event is similarly filtered using a low-dimensional set of temporal kernels. Each individual neuron’s firing rate at each time bin is a nonlinear function (here, *f* (·) = exp(·)) of the sum of a linear weighting of the low-dimensional stimulus subspace, a linear weighting of the touch-bar subspace, and recent spike history. Spiking is given as a Poisson process given the instantaneous rate. Because we do not include interactions between neurons here, this model can be applied to a set of single-neuron recordings. However, the model can readily be extended to include interactions in simultaneously recorded populations (Pillow et al., 2008; Pandit et al., 2020). (**B**) The model represents the different stimulus directions and categories — including whether it is the sample or test stimulus — as vectors in a low dimensional space where the dimensionality is the number of factors. The vectors change over time given the temporal kernels. The models we focus on here constrain the direction tuning, but not category, to be constant over the sample and test stimuli. The full GMLM allows for flexible direction tuning (i.e., each stimulus direction is a distinct point) and the cosine-tuned GMLM constrains the direction tuning to an ellipse. Thus, dimensionality reduction in the GMLM can take into account that temporal dynamics and stimulus tuning information can be shared across the two stimulus presentations. Individual neurons’ stimulus tuning is a linear projection of the low-dimensional space.

By applying the GMLM to the LIP populations, we quantified population-level differences in LIP activity between animals and compared those differences with behavioral performance with respect to the animals’ training histories. We found category and direction selectivity in LIP during the DMC task in all subjects. However, we found stronger cosine-like motion direction tuning in LIP during the DMC task in monkeys first trained on the DMS task compared to monkeys trained only on categorization. During the test stimulus presentation when the monkeys had to compare the incoming test stimulus to the remembered sample stimulus, sample category could be more reliably decoded from LIP responses irrespective of the test-stimulus direction in the DMC-only monkeys than in the pretrained monkeys. Behaviorally, the pretrained monkeys were more likely to make categorization errors when the sample and test stimuli were similar than the DMC-monkeys. Additionally, we introduce dynamic spike history within the GMLM which reveals a difference in oscillatory, single-trial dynamics during the delay period between the DMC-only and pretrained monkeys. Together these results suggest that different subjects may recruit distinct behavioral and neuronal strategies for performing the DMC task, and that long-term training history may play a role in shaping these differences. Low-dimensional encoding of the DMC task in LIP may therefore reflect training history or a particular task strategy (or both).

## 2 Results

### 2.1 Dimensionality reduction in LIP with the GMLM

We aimed to describe how task-related responses in LIP are shared across neurons in a population by reducing the dimensionality of the eight LIP populations using GMLMs. The GMLM uses a low-dimensional set of components to describe the population responses during each trial as a multilinear function of the task variables (Fig. S1). To place the DMC task into the GMLM framework, the motion stimuli were linearized as two sets of regressors: (1) timing events to indicate the onset of a stimulus (sample or test) and (2) stimulus identity regressors that encode direction, category, or if the stimulus is the sample or test. The model’s parameters include a set of stimulus components, where each component contains a single temporal kernel (or linear filter) and a set of weights for the stimulus identity. Each component temporally filters the stimulus onset events, and weights the filtered stimuli linearly by stimulus identity. As a result, each individual component contributes to the population encoding for all stimuli (not just a single motion direction or sample/test presentation). Each individual neuron’s tuning to the motion stimuli is a linear combination of the stimulus components. The model also includes a low-dimensional set of components to represent the touch-bar release event: a set of temporal kernels describe the population response to a touch-bar release such that each neuron’s touch-bar tuning is a linear combination of those kernels. Each spike train is then defined as a Poisson process in which the instantaneous firing rate is given by the sum of the filtered stimuli and touch-bar release, plus a linear function of recent spike history. The set of stimulus components is a low-rank tensor that represents population tuning to the motion stimuli in a low-dimensional subspace which captures shared response dynamics across neurons in the population. The factorized representation of the stimulus into temporal and identity weights captures shared temporal dynamics between different directions or between sample and test stimuli. As the number of components (i.e., rank) in the kernel tensor increases, the model approaches a GLM fit to each cell individually.

We designed a set of four nested models (i.e., different linearizations of the motion stimuli with increasing complexity) in order to assess what stimulus information is encoded by an LIP population (Fig. S1E-I). The simplest model was the no category or direction tuning model. In this model, the linear weights for the stimulus identity only defined whether the stimulus was the sample or test. This model can only capture the average trajectories in time during the task over all stimulus directions. The second model, the category only model, includes stimulus category weights, but does not consider specific motion direction. The category only model includes stimulus identity information for the category one and category two motion directions for both the sample and test stimuli. This way, the model can capture category tuning, which may be different for the two stimulus presentations. While the DMS task had no category, we still fit category weights to the DMS populations as a control (i.e., to ask what the model produces if category was not actually a behavioral factor in the task). The third model included cosine direction tuning and category. In addition to the category weights from the previous model, this GMLM included two coefficients for the sine and cosine of the motion direction. The cosine and sine weights were the same for both sample and test stimuli. Thus, this model constrains the geometry of direction tuning to lie on an ellipse in the low-dimensional space (Fig. 2B). The final model, the full model, extends the cosine direction tuning model by allowing different weights for each individual motion direction, rather than constraining the direction information to be cosine tuned.

The full GMLM fit to the LIP population from monkey B is shown in Fig. 3A,B. The model has seven stimulus tensor components, each with a temporal kernel (left column). The temporal kernel is scaled for each of the six directions in the task to give a temporal kernel for the sample stimulus (middle column). Next, we include additional test category kernels, which are added to the direction kernels in the middle column, to describe the response to the test stimulus (right column). Different components can have different temporal response dynamics and different stimulus tuning properties: for example, component 5 shows strong differentiation between the two stimulus categories (red and blue), while component 2 does not. Each cell’s response tuning (linear kernels) is then a linear combination of the components. The tuning to the sample directions for individual cells is illustrated in Fig. 3C,D. The GLM fits to the individual cell and corresponding PSTH are shown below the GMLM fit for comparison. We note the total firing rates fit by the models are a combination of the stimulus filters, baseline rate, spike history effects, and firing rate nonlinearity. As a result, the PSTHs and filters do not match exactly.

**Figure 3:**
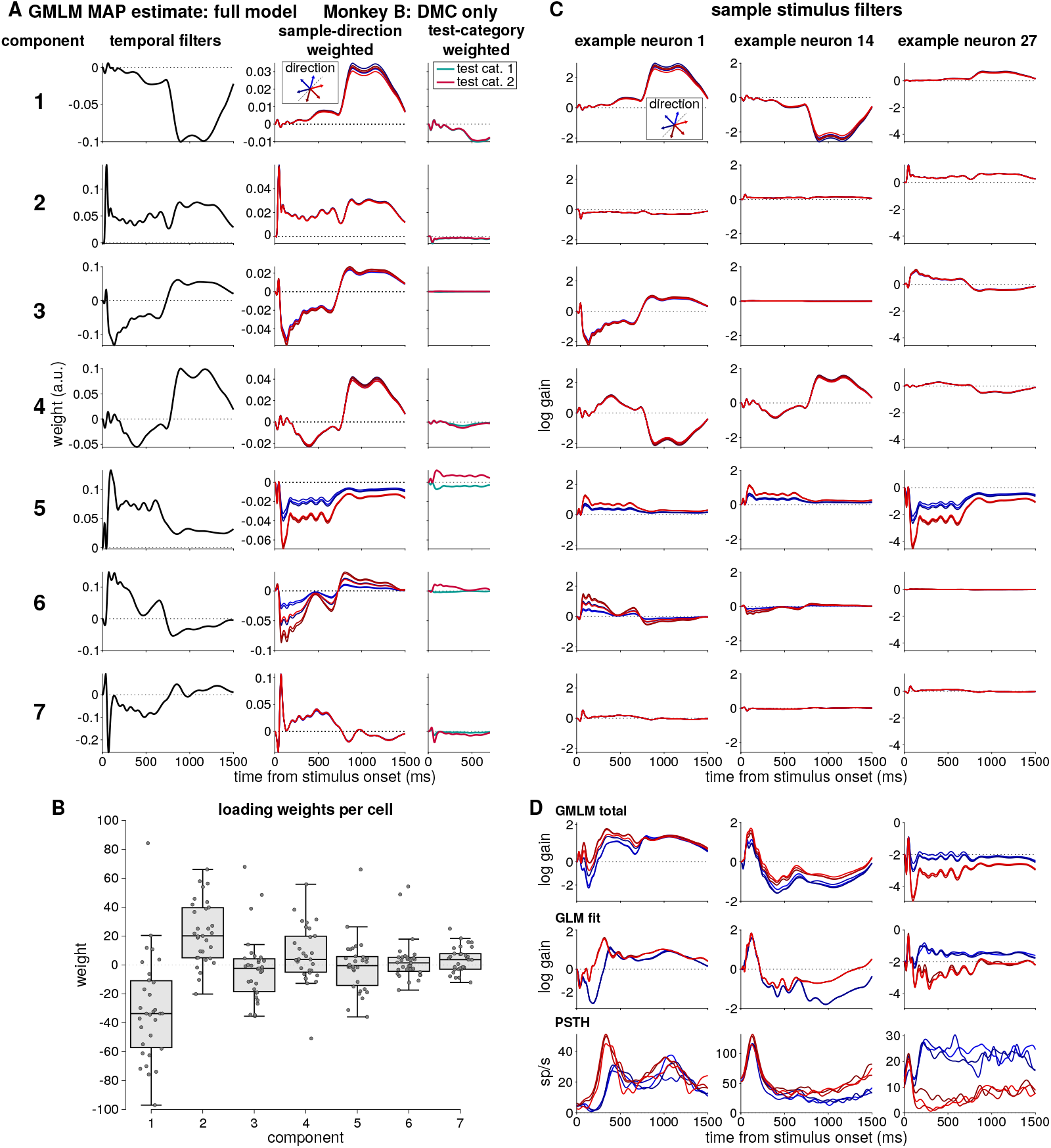
GMLM with seven stimulus components fit to the LIP neurons recorded from monkey B during the DMC task. (**A**) The seven GMLM stimulus components that define the population responses to the motion stimuli are given in each row. (left) The temporal kernels for each component (normalized). (middle) The temporal kernels weighted for each sample stimulus direction. (right) The temporal kernels weighted by the test category (the test stimulus kernels are shown shorter than the sample stimulus kernels because trial ends after the test stimulus presentation). The total kernels for each test direction are computed by adding the test category kernels to the sample direction filter. As a result, the sample and test kernels have the same direction tuning, but the kernels may have different category tuning (e.g., component 5). (**B**) The distributions of weights for each component across all neurons. Each dots is the weight for one neuron for the given component. (**C**) Each column shows the model fit of the sample stimulus kernels for three example neurons. The first seven rows show the seven components scaled by each neuron’s linear weighting of the component (the middle column of part A with neuron-dependent scaling from B). (**D**) Total sample stimulus tuning for the three example cells. The total GMLM stimulus kernels (i.e., the sum of rows in C; top row), the GLM fit to the individual cell (middle), and the PSTH conditioned on sample stimulus direction for comparison (bottom). The low-dimensional set of kernels for the touch-bar release are shown in Fig. S2.

The parameterization in the cosine-tuning and full models assumes the direction tuning (but not category tuning) is the same for both sample and test stimuli: that is, the difference between the kernels for two motion directions within the same category are the same for both the sample and test stimulus. Such direction tuning constancy would be consistent with common bottom-up, direction-tuned input from sensory areas such as MT for the two stimulus presentations. The model still includes test category filters, which allow for different category tuning or direction-independent gain differences between sample and test stimuli. We found that including separate sample-test direction tuning in either model did not improve the model fit (Fig. S3A). Additionally, comparing sample and test direction weighting in the low-dimensional GMLM component space showed similar direction preferences for the two stimuli (Fig. S3B). Thus, the GMLM framework can both constrain the parameters to enforce constant direction tuning between the two stimuli, and test statistically whether that assumption holds.

### 2.2 Model selection and dimensionality

We first determined the dimensionality of stimulus-related activity in the LIP population responses. We varied the number of components to include in the GMLM (i.e., the rank of the stimulus kernel tensor). We compared the full GMLM to the corresponding single-cell GLM fits, where the GLMs represent the “full-rank” model. We computed the average likelihood per trial averaged over the neurons in each population for the GMLM fit, relative to the GMLM without any stimulus terms (Fig. 4A). We selected the rank by the number of components needed to explain, on average, 90 % of the explainable log likelihood per trial over all the neurons in each population (Fig. 4B). The GMLM required 7 to 12 stimulus components per population, thereby using only a fraction — less than 8 % — of the number of parameters compared to GLM fits to individual cells (Fig. 4C).

**Figure 4:**
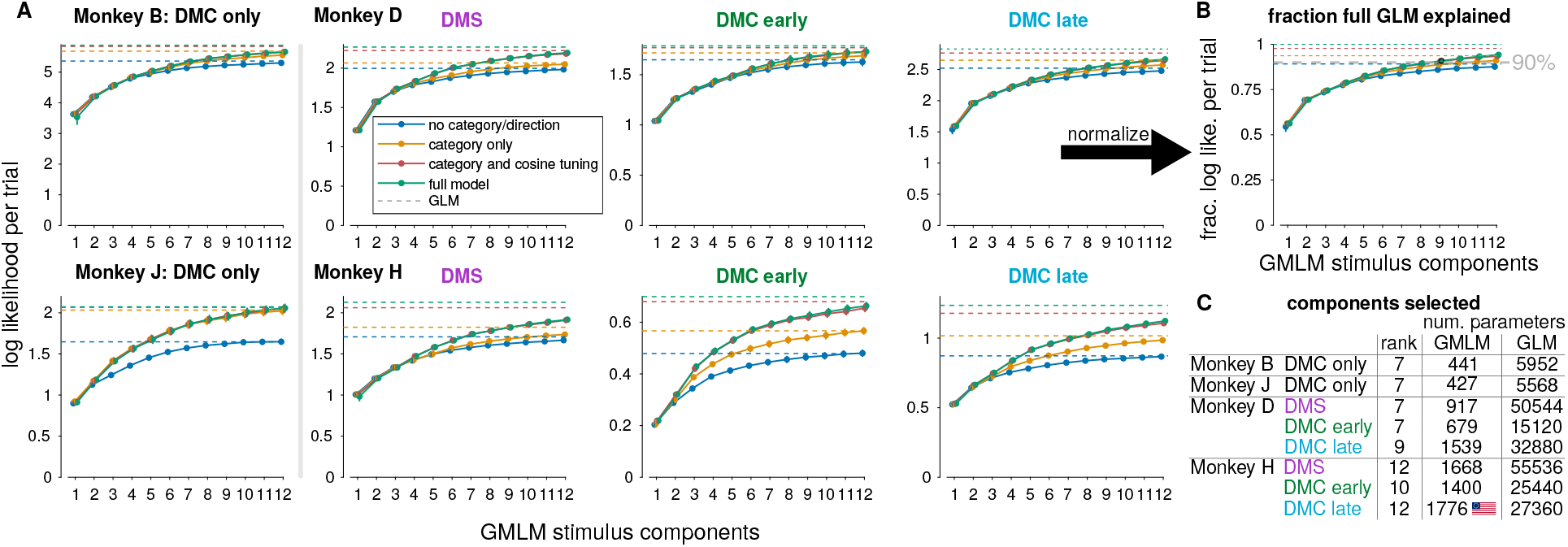
Rank selection in the GMLM. (**A**) The mean cross-validated log likelihood per trial averaged across neurons of the GMLM as a function of the stimulus kernel tensor rank relative to the model without any stimulus terms (i.e., the rank 0 model) for each of the eight LIP populations. Each trace shows the log likelihood for a single model parameterization: no stimulus information (i.e., only mean temporal dynamics; blue), category only (no specific direction information; yellow), category plus cosine direction tuning (red), and the full model with flexible direction tuning (green). The dashed lines show the mean cross-validated likelihood of the GLMs, which correspond to the full-rank model. (**B**) The fraction of log likelihood of the full GLM (fraction of the log likelihood that could be captured by the GMLM) was used to select the GMLM rank for each LIP population, shown here for monkey D, DMC late. This fraction is the cross-validated log likelihood divided by the log likelihood of the full GLM (dashed green line), and the threshold for rank selection was 90 % of the log likelihood per trial. (**C**) Number of stimulus components (rank) selected for the GMLM for each LIP population. The number of stimulus parameters in the low-rank GMLM (full model) is compared to total parameters in the equivalent single-cell GLM fits for all cells in each population.

We then asked how much each task variable contributed to the low-dimensional LIP responses. To do so, we quantified model fit as we monotonically increased the complexity in the nested models by including more information about the stimulus identity and category. A majority of the log likelihood was accounted for by the GMLM without category or direction tuning in all populations, which is consistent with many previous dimensionality reduction results (Kobak et al., 2016). The category-only GMLM captured a greater percentage of the log likelihood over the no-category model in the DMC late populations than in the DMS populations (Monkey B, DMC only 2.5 ± 0.3 %; Monkey J, DMC only 13.8 ± 0.9 %; Monkey D, DMS 2.1 ± 0.3 %, DMC early 1.4 ± 0.7 %, DMC late 2.8 ± 0.2 %; Monkey H, DMS 3.3 ± 0.2 %, DMC early 11.3 ± 0.7 %, DMC late 9.5 ± 0.3 %; mean percentage cross-validated log likelihood accounted for by the category-only GMLM minus the no-category GMLM where the errors are ±2 SEM over the cross-validation folds).

Cosine direction tuning during the DMC task accounted for a larger improvement of the model fit over the category only model for the pretrained monkeys than the DMC-only monkeys. (monkey B, DMC only 1.0 ± 0.4 %; monkey J, DMC only 0.2 ± 0.7 %; monkey D, DMS 4.1 ± 0.3 %, DMC early 1.6 ± 1.0 %, DMC late 1.9 ± 0.1 %; monkey H, DMS 8.2 ± 0.6 %, DMC early 11.9 ± 0.5 %, DMC late 9.9 ± 0.6 %; mean percentage cross-validated log likelihood accounted for by the cosine-tuned GMLM minus the category-only GMLM). Thus, direction-tuning played a stronger role in the pretrained monkeys’ LIP activity than in the DMC-only monkeys.

We tested the adequacy of cosine parameterization of direction tuning by comparing the more flexible full model. The cosine direction tuning model was comparable to the full model for all populations (monkey B, DMC only 0.2 ± 0.2 %; monkey J, DMC only 0.3 ± 0.7 %; monkey D, DMS 0.2 ± 0.2 %, DMC early 0.2 ± 0.1 %, DMC late 0.2 ± 0.2 %; monkey H, DMS 0.3 ± 0.4 %, DMC early 1.1 ± 0.5 %, DMC late 1.3 ± 0.3 %; mean percentage cross-validated log likelihood accounted for by the full GMLM minus the cosine-tuned GMLM). For these tasks, the direction tuning in the population could therefore be approximated as an ellipse (and thus embedded within a plane).

### 2.3 Low-dimensional response to the sample stimulus

To gain intuition about how LIP dynamics may support the transformation of motion direction input into a representation of category, we visualized the low-dimensional population tuning to the sample stimuli. The top three dimensions of the trajectories show large, stimulus-independent transient responses (Fig. 5, inset; Fig. S4). This is consistent with the large fraction of the data explained by the GMLM without category or direction tuning. We therefore subtracted the mean response over stimuli and plotted the top three dimensions in the mean-removed responses (Seely et al., 2016). The two DMC-only LIP populations showed primarily two-dimensional responses with strong category separation (Fig. 5). For the pretrained monkeys, the trajectories during the DMS task reflected the stimulus geometry: the model shows two-dimensional transient activity organized by stimulus angle. The responses show little stimulus-specific persistent activity during the delay period (Sarma et al., 2016): the trajectories return to the origin after stimulus offset. During the DMC early phase, the low-dimensional LIP response reflects the stimulus geometry, but the top dimension is aligned to the task axis (i.e., blue and red directions are separated along dimension 1). The trajectories are still two-dimensional without clear delay period encoding. After training was completed, monkey D’s DMC late LIP activity showed strong direction tuning during the stimulus presentation which is elongated along the task axis (that is, the axis most oriented to category along the 135°-315°stimulus directions). In contrast, LIP in monkey H had a three-dimensional stimulus response in the late period: two dimensions reflecting the circular motion directions during the stimulus presentation and a third orthogonal axis for category that was sustained through the delay period. Similar orthogonal stimulus input and working memory representations have been observed in other decision making tasks (Aoi et al., 2020; Libby & Buschman, 2021). In summary, the low-dimensional stimulus components of the LIP activity differed across ani-mals such that the pretrained monkeys’ LIP showed strong, circular representations of motion direction, while the DMC-only monkeys had lower-dimensional responses that more strongly reflected category.

**Figure 5:**
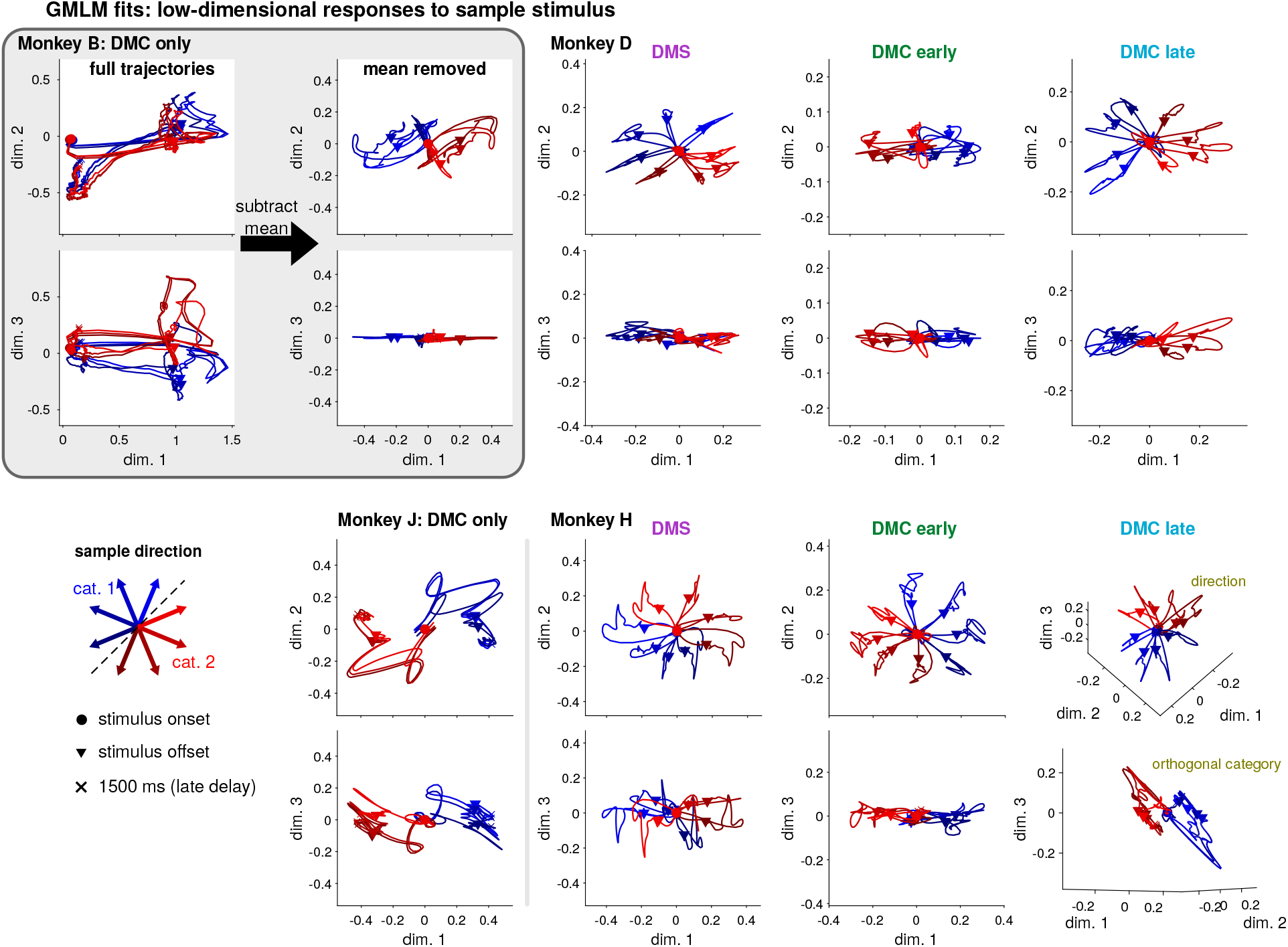
Low-dimensional representations of motion direction and category during the first 1500 ms of each trial (sample stimulus presentation and delay period). The top three dimensions of the GMLM’s sample stimulus encoding for each of the eight LIP populations with the mean response over all directions removed. The inset for monkey B shows the top three dimensions including the mean (left) and the top three dimensions that remain after removing the mean (right). The two plots for each monkey show the top dimension on the x-axis plotted against the second or third dimensions on the y-axes (except for monkey H shown in the 3-D plots). The red and blue traces show the response to each motion direction from stimulus onset (circles), to stimulus offset (triangles), and into the delay period (x’s denote 1500 ms after sample motion onset). The three-dimensional trajectories (of the cosine-tuned model) are shown as a function of time in Fig. S5.

### 2.4 Quantifying the geometry of category and direction in the stimulus subspace

To go beyond visualization of the low-dimensional subspace, we wished to quantitatively assess the geometry of category- and direction-dependent responses in LIP. Here, we focused on the cosine-tuned GMLM. Given this choice of parameterization, direction and category could be decoupled, while still capturing a similar subspace to the full model (Fig. S5). At each point in time, category was encoded along a vector while direction (parameterized by angle) was encoded on an ellipse in the stimulus subspace (Fig. 6A). The ellipse could be circular, which would represent motion directions uniformly, or elongated so that the population representations are biased towards a preferred motion direction. We compared the norm of major and minor axes of the direction ellipse and the norm of the sample category vector (Fig. 6B) as a function of time relative to stimulus onset. We conducted a Bayesian analysis of the GMLM’s sample stimulus subspace to take into account uncertainty in the model fit given the posterior distribution of the model parameters.

**Figure 6:**
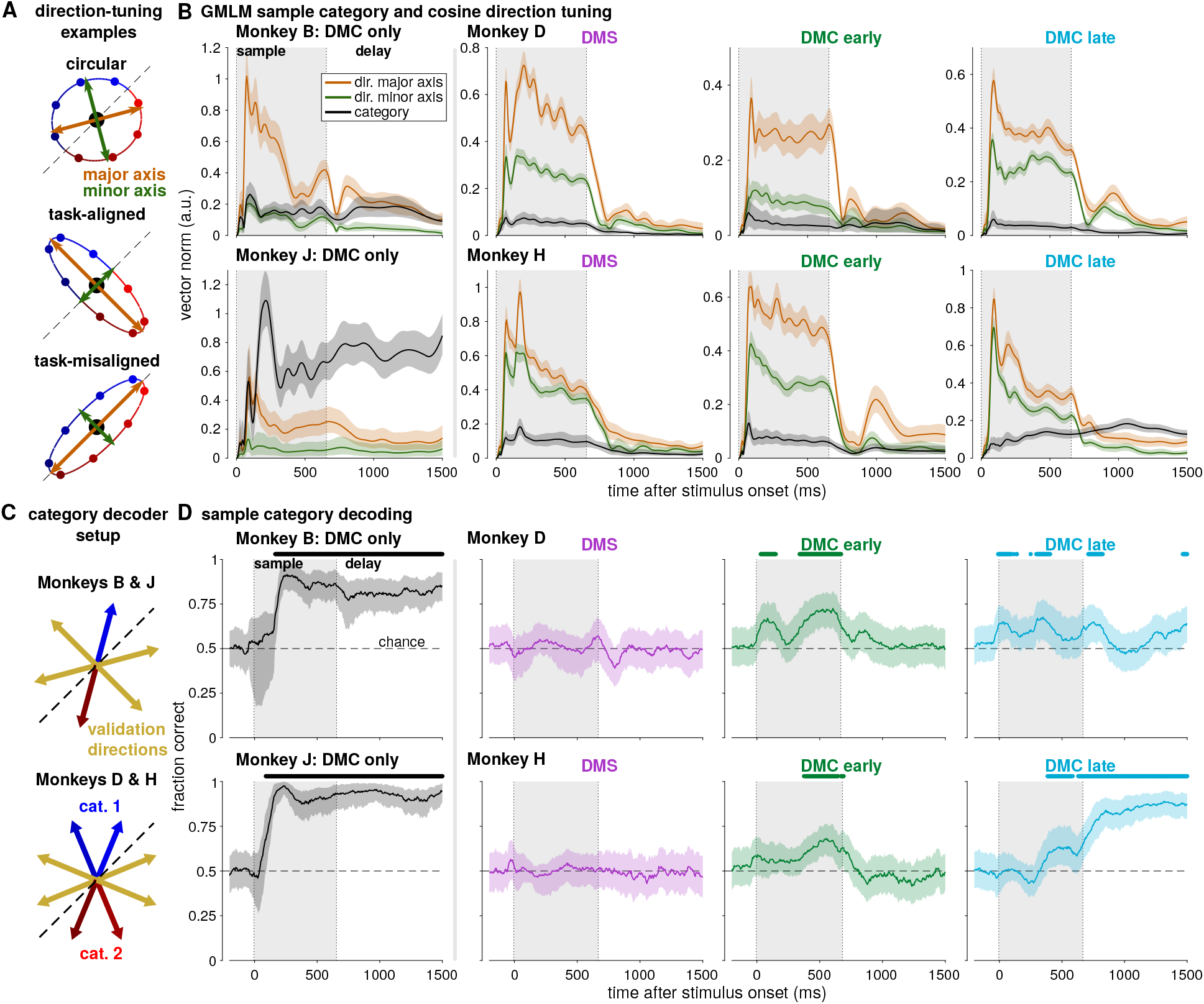
Quantification of category and direction encoding in LIP. **A** Diagram of direction encoding in the cosine-tuned GMLM. Motion stimulus direction is encoded as an ellipse in the low-dimensional stimulus space. The ellipse has major (orange arrows) and minor (green arrows) axes which define its shape. If the axes are of similar length, tuning is approximately circular and the motion directions are evenly distributed in the low-dimensional space (top). If the major axis is elongate compared to the minor axis, the population shows a preferred direction which may be aligned with category (middle) or the null direction (bottom) or anywhere in between. Category is encoded as a vector in addition to the direction ellipse, and the category vector is constant for all motion directions within a category (in contrast to the task-aligned direction tuning which places near-boundary directions closer together). **B** Bayesian estimate of the geometry of the sample stimulus tuning for the eight LIP populations. Each plot shows the norm of the sample category vector (black) and the norms of the major (orange) and minor (green) axes of the direction tuning ellipse for an LIP population as a function of time relative to the sample stimulus onset. The solid lines denote the posterior median at each time point, and the shaded regions denote 99 % credible intervals. If the major and minor axes have equal norms, then direction would follow a circle in the low-dimensional space. **C** The training and test scheme for a direction-independent category decoder. Psuedopopulation trials were created from 50 random trials sampled with replacement for each direction from each cell. For monkeys B and J, the decoder is trained using one direction from each category (180° apart; red and blue) and validated on the remaining four directions (yellow). For monkeys D and H, the decoder is trained using two adjacent directions from each category (again using opposite directions from each category; red and blue) and validated on the remaining four directions (yellow). **D** Median decoder generalization performance as a function of time for each of the eight LIP populations. The decoder was trained and tested using spike counts in a 200 ms window centered at the time relative to sample stimulus onset on the x-axis. The shaded regions denote a 99 % confidence interval over 1000 random pseudopopulations. Symbols denote decoding significantly greater than chance (50 %; *p* < 0.01 Benjamini-Hochberg corrected, one-sided bootstrap test).

The DMC-only LIP population subspaces showed strong category tuning relative to direction tuning. The category vector in monkey B was of similar magnitude to the minor axis of the direction ellipse during stimulus presentation, and stronger during the delay period. The category vector in monkey J was larger than the direction tuning ellipse throughout the stimulus presentation and delay period. The direction tuning ellipses were elongated along a particular motion direction, rather than circular. Additionally, the direction ellipse aligned both with category and with the choice biases in monkeys B and J on trials where the sample motion direction was ambiguous (Fig. S6). The sample stimuli on ambiguous trials were placed on the category boundary, and the monkeys were rewarded randomly. The ambiguous trials were not used to fit the GMLM. Thus, the stimulus components in the DMC-only populations reflected category-specific input selection.

In the pretrained monkeys, the DMS populations showed strong direction tuning, which in monkey H was nearly circular or uniform (i.e., the major and minor axes of the direction ellipse were of similar length). During the late DMC sessions — but not during the DMS task — monkey D’s direction ellipse was aligned with the task category (i.e., the major axis was along the 135°-315°angles; Fig. S6C). The same task-aligned direction encoding during the DMC task was not observed in monkey H. In both the monkey D late DMC and monkey H early DMC populations, we found stimulus offset activity in the direction ellipse, but not in the category vector. As a result, individual neurons may appear to respond more strongly for a particular category early in the delay period, but the model accounted for this as a direction-tuned response rather than category-specific encoding. The monkey H early and monkey D early and late DMC populations did not have large category vectors, and the low-dimensional activity instead reflects an elongated direction tuning ellipse (i.e., the major axis is larger than the minor axis) during stimulus presentation. In the monkey H late DMC population, we observed a slow increase in the category vector length over time in the trial, which does not surpass the magnitude of direction tuning until the delay period.

We then asked how the subspace geometry could affect how decoding methods assess category selectivity in LIP. We applied a linear decoding technique previously proposed to reveal category representations independent of motion direction (Swaminathan et al., 2013; Sarma et al., 2016). The decoder classifies the sample category based on spike counts from pseudopopulation trials. We trained and validated the decoder on trials from orthogonal sets of stimulus directions (Fig. 6C). The logic of the decoder is that, if motion direction is represented in the population circularly without any additional category-specific responses, the decoder will not generalize across the training and validation conditions. The DMS populations provided a control for the method, because the monkey was not yet trained to classify motion category. We indeed found no significant category decoding in the two DMS populations (Fig. 6D).

The decoder performances during the DMC task were consistent with the GMLM stimulus subspaces. Category could be decoded with high accuracy in monkeys B and J early in the stimulus presentation and throughout the delay period (Fig. 6D). Similarly, the GMLM analysis had found strong category tuning beginning early in the stimulus period and continuing through the delay in those populations. The results were different in the pretrained animals. In both DMC early and monkey D’s DMC late populations, we found decoding above chance during stimulus presentation, but not during the delay. The task-aligned, non-circular response to stimulus direction in monkey H DMC early and monkey D DMC early and late enabled the decoder to generalize across conditions due to over-representation of signal along the task dimension (135°-315°), rather than a category vector independent of direction. In contrast, in the DMC late population for monkey H, the decoder only found weak decoding late in the sample stimulus presentation, which became strong during delay period. The orthogonal category dimension of monkey H DMC late is only stronger than the circular direction coding during the delay period, corresponding to the onset of significant category decoding. The decoder’s failure to generalize during the early sample period can therefore be explained by strong direction selectivity swamping the weaker, orthogonal category signal.

### 2.5 Comparing the sample and test stimuli

The DMC task requires different computations for the test and sample stimuli: the category of the sample stimulus must be computed and stored in short-term memory, while the test stimulus must be compared to the stored sample category. Recent work has suggested that LIP linearly integrates the test and sample stimuli during the test period of the DMC task while prefrontal cortex shows more nonlinear match/non-match selectivity (Zhou et al., 2021). We therefore compared the LIP responses to the test stimulus to the population responses to the sample stimulus. In the GMLM fits to the DMC populations, we found that test category tuning during the test stimulus presentation was weaker than sample category tuning during the sample presentation (Fig. S7). This can be seen in the low-dimensional subspace for monkey H (Fig. 7A). During the sample stimulus, the subspace reflected category tuning orthogonal to the motion direction response (Fig. 5, bottom right). However, we did not find the same category-selective response to the test stimulus in the stimulus subspace. Addition ally, LIP population activity projected onto the touch-bar (motor response) subspace showed strong match/non-match separation with little category selectivity (Fig. S8). Thus, LIP does not appear simply to extract and sum the categories of the two stimuli to compute match or non-match.

**Figure 7:**
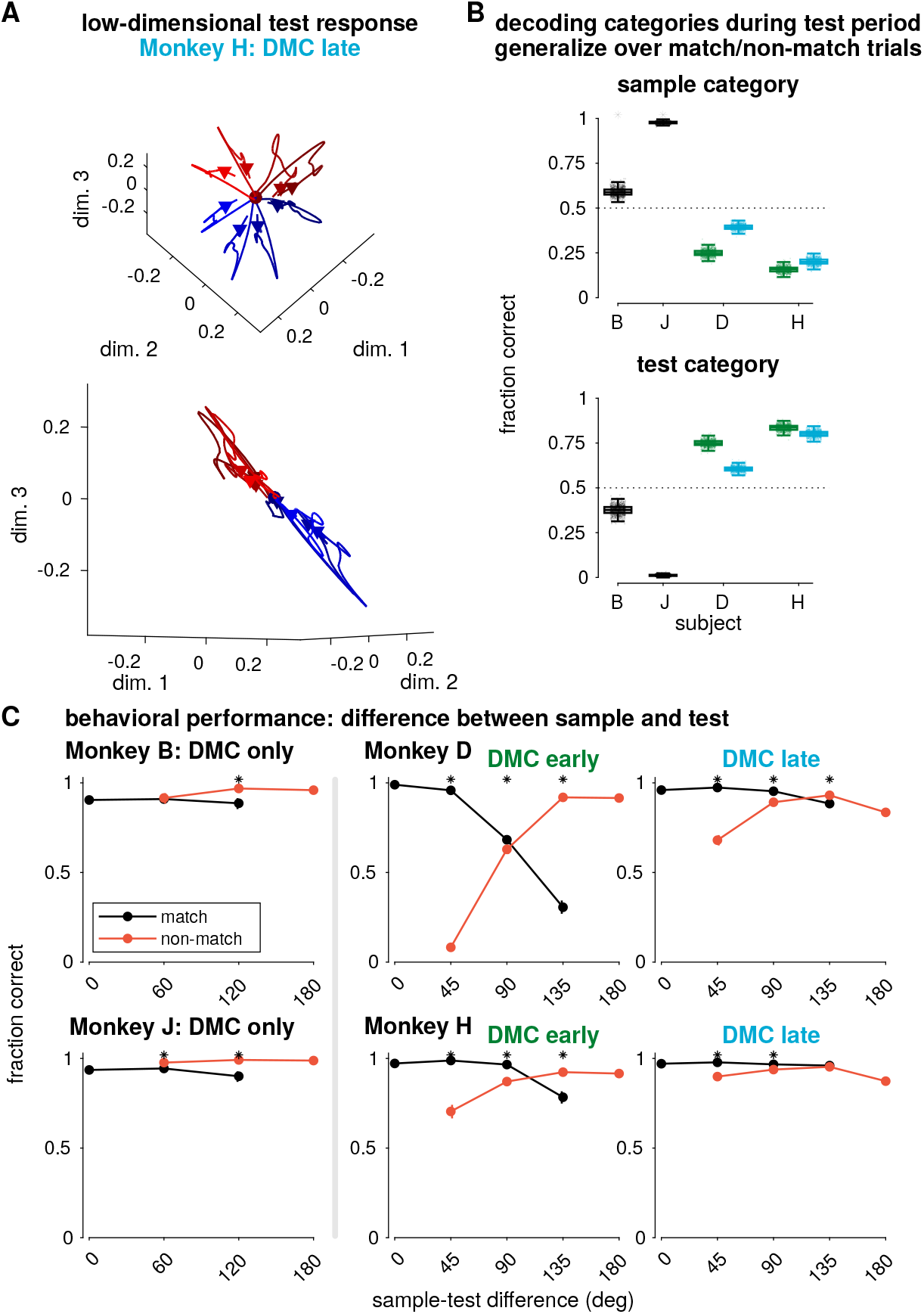
Matching the test stimulus to the stored sample in the DMC task. **A** The low-dimensional test stimulus response for each direction for monkey H, late DMC with the mean response removed projected into the same dimensions as in Fig. 5 (bottom right). **B** Decoding accuracy of sample (top) or test (bottom) category using the spike counts during the first 200 ms of the test stimulus (excluding the motor response for 95.7 % of match trials). The decoder was trained on trials from all stimulus directions, but only from match (or non-match) trials and then tested on non-match (or match) trials. All decoders generalized significantly different than chance (50 %; *p* < 0.01 Benjamini-Hochberg corrected, two-sided bootstrap test). **C** Average performance as a function of the difference in angle between the sample and test stimulus, sorted by match/non-match trials in the DMC task (error bars show a 99 % credible interval). Asterisks indicate match and non-match are significantly different (*p* < 0.01, two-sided rank sum test, Benjamini-Hochberg corrected).

We tested if the stored sample category and incoming test stimulus category were separable in the LIP population responses during the test stimulus presentation. We used linear classifiers to decode the sample or test category from pseudopopulation spike counts during the first 200 ms of the test stimulus presentation. The decoders were trained on trials of all stimulus directions. However, the training set consisted of only match (or non-match) trials, while the validation set included only non-match (or match trials). For the two DMC-only animals, monkeys B and J, the sample category decoder generalized across the two conditions (i.e., performed better than chance). The test category decoder, however, performed much worse than chance (Fig. 7B). Thus, the decoding axis for test category switched signs across match and non-match trials. We observed the opposite pattern in the pretrained monkeys: the decoders generalized to classify the test stimulus category, but not the sample. In the DMC-only animals, sample category can therefore be read out by a single linear decoder regardless of the test stimulus identity, which is consistent with stronger separability of the remembered sample category and the incoming test stimulus in the DMC-only monkeys than in the pretrained monkeys. Increased separability suggests a coding scheme that reduces interference between the stored sample stimulus category and the specific test stimulus direction (Libby & Buschman, 2021).

We then asked how the monkeys’ performances depended on the similarity between the sample and test stimuli. We compared the monkeys’ accuracy as a function of distance between test and sample directions (Fig. 7C). The pretrained monkeys showed a different pattern of accuracy than the DMC-only monkeys. At small sample-test differences, the pretrained animals showed better performance on match than non-match trials while the DMC-only monkeys perform similarly or better on non-match. Additionally, the pretrained animals showed greater dependence on distance. These effects were greatest during the early DMC training phase, but they persisted after extensive training on the order of several months (the total number DMC training sessions between the DMC early and DMC late periods was 78 for monkey D and 65 for monkey H). Therefore, stimulus similarity — which was relevant in the DMS task— affects categorization behavior more strongly in the pretrained monkeys than the DMC-monkeys, reflecting the monkeys’ strategy.

### 2.6 Single-trial dynamics during the delay period

Neural dynamics during single-trials may reflect aspects of sensory processing and working memory beyond the mean firing rate (Fontanini & Katz, 2008; Miller et al., 2018). For example, working memory may be supported by persistent activity (Constantinidis et al., 2018) or oscillatory bursts (Lundqvist et al., 2016) while stimulus-related activity may exhibit strong transient responses with quenched variability (Churchland et al., 2010). We therefore sought to characterize non-Poisson variability in single trials in LIP during the DMS and DMC tasks, which could reflect signatures of different strategies in performing the tasks. The GLM framework accounts for non-Poisson variability or single-trial dynamics by conditioning firing rates on recent spiking activity through a spike history filter, an autoregressive term which can reflect a combination of intrinsic (e.g., refractory periods) and network properties (e.g., oscillations) (Truccolo et al., 2005; Weber & Pillow, 2017; Zoltowski et al., 2019). Typically, the GLM assumes the spike history filter to be constant: that is, spike history has the same effect on spike rate throughout a trial. While fixed spike history effects may be an appropriate assumption in early sensory regions under stimulation with steady-state stimulus statistics, spiking dynamics in LIP might vary between the stimulus presentation and the delay period due to the transition in behavioral task demands between these two periods of the task (Hart & Huk, 2020). In order to quantify spike history effects in the DMS and DMC tasks, we extended the GMLM to include *dynamic spike history filters* which allows the autoregressive dynamics to change throughout a trial (Fig. 8A). The dynamic spike history in the GMLM was a low-rank tensor with a linear spike history component and a gain term relative to the stimulus timing (Harris et al., 2019). As a result, the model learns how each neuron’s spike history filter changes during the course of a trial relative to task events, and can therefore capture distinct dynamics between stimulus-driven and delay periods of the trial.

**Figure 8:**
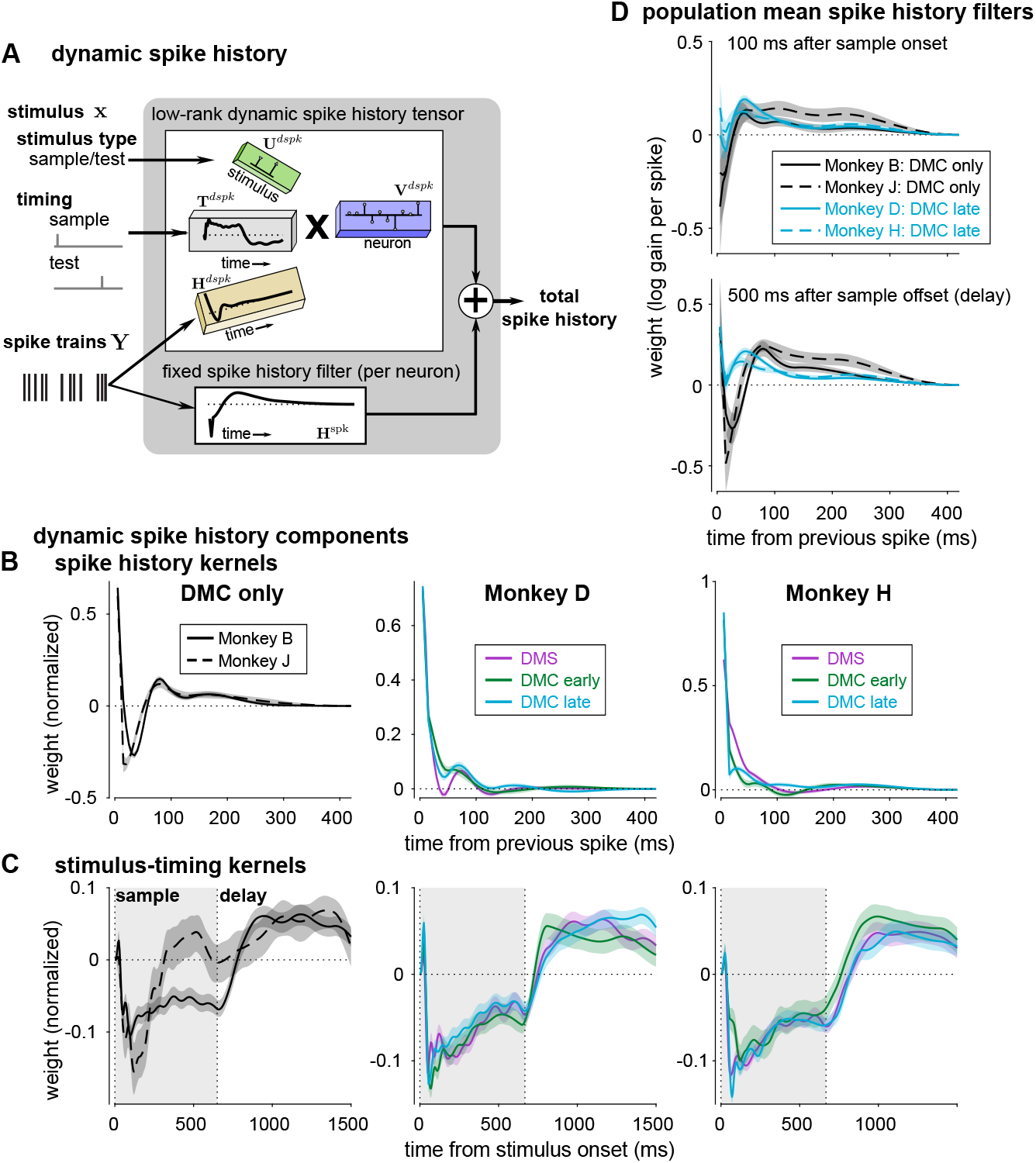
Dynamic spike history captures distinct stimulus-driven and delay-period dynamics. **A** The dynamic spike history filter is modeled as a low-rank, four-way tensor. The tensor includes two temporal kernels: one which filters spike history (gold) and a second which determines the weighting of the spike history component relative to the stimuli (gray). The spike history is scaled by stimulus identity (for simplicity, limited to sample or test stimulus weights only, without any category information). Each neuron adds the (weighted) dynamically filtered spike history to the neuron’s constant spike history filter. The total spike history at any point in a trial is still a linear function of past spiking activity, but the effective linear kernel can change during a trial. The normalized rank-1 dynamic spike history components: **B** dynamic spike history kernels and **C** stimulus-timing kernels for each population (posterior median and 99 % credible interval). The left columns shows the two DMC-only monkeys, and the middle and right columns show all three training stages for monkeys D and H respectively. **D** The population mean effective spike history filters at two points in the task given the rank-1 dynamic spike history for the four fully trained DMC populations (mean of MAP estimate of filters ±2 SEM). (top) The population mean spike history at 100 ms after sample stimulus onset. (bottom) The population mean spike history during the delay period (500 ms after sample stimulus offset). Positive weights indicate that a previous spike at the given lag increases a neuron’s probability of firing, while negative weights indicate that spiking is suppressed.

We fit the GMLM with a single dynamic spike history component to the LIP populations. Including dynamic spike history improved the cross-validated model performance for all populations (Fig. S9A). The GMLM found similar dynamic spike history kernels for the two DMC-only monkeys, which showed oscillatory dynamics at approximately 12 to 14 Hz (low-beta; Fig. 8B). In contrast, the dynamic spike history kernel for the pretrained monkeys at all training stages was dominated by a faster timescale decay (time constants monkey D 10.2, 14.8 and 11.8 ms and monkey H 22.5, 9.0 and 6.6 ms DMS, DMC early, DMC late respectively), which suggests stronger gamma-frequency bursts. The stimulus-timing kernels showed that the dynamic spike history generally aligned with stimulus onset and offset (Fig. 8C). One notable exception was monkey J: the timing showed only a short transient gain after stimulus onset. The timing of this kernel corresponded with the strong category-dependent transient response in monkey J (Fig. 6B), and thus raises the possibility that the network enters a memory-storage state prior to stimulus offset.

Lastly, we examined population differences in the total spike history: the dynamic spike history filter (which depends on time in the trial) plus the individual neurons’ fixed spike history filters. We computed the population mean spike history kernel at two different points (Fig. 8D): during stimulus-driven activity (100 ms after sample stimulus onset) and during the delay period (500 ms after sample offset). The mean spike history in the DMC-only monkeys showed a pronounced oscillatory-like trough during the delay period, compared to the pretrained monkeys (Fig. 8D bottom). Spike history differences between the populations were less evident during stimulus presentation. These results were consistent for higher-rank dynamic spike history tensors (Fig. S9). Thus, the structure of single-trial variability during the delay period differed between DMC-only and pretrained monkeys, but was similar within each pair, which suggests that the balance of beta- and gamma-frequency driven activity during the delay period differed between the animal pairs.

## 3 Discussion

Here we examined the low-dimensional geometry of task-related responses during a motion-direction categorization task in LIP in two pairs of monkeys performing the same motion categorization task, but with different training histories. In the monkeys that were pretrained on a motion-discrimination DMS task, we found similar direction-dependent activity in LIP activity during the DMS and DMC task: two-dimensional direction-encoding subspaces that reflected the stimulus geometry. Moreover, uniform direction tuning remained a dominant feature of this subspace after training on the DMC task. The common direction tuning observed across the sample and test stimuli could reflect cosine-like signals from sensory regions such as the area MT (Born & Bradley, 2005; Jazayeri & Movshon, 2006; Fanini & Assad, 2009). In contrast, the monkeys trained only on the categorization task showed stronger category tuning and category-aligned direction tuning in LIP compared to activity in animals first trained on the DMS task. Performing the categorization task may involve computations including input selection and local and/or top-down recurrent dynamics. Our findings indicate that differences in the sequences of tasks learned by the animals over long periods may result in different network configurations that perform the same task, perhaps manifesting in different behavioral strategies.

We hypothesize that these differences may be indicative of the pretrained monkeys still using computational strategies learned for the DMS task. Indeed, the pretrained monkeys’ behavior showed greater dependence on the angular difference between sample and test stimuli than DMC-only monkeys, which was a key factor in solving the DMS task. Because the tasks used the same stimuli and shared many of the same correct or incorrect sample-test pairings, the same neural machinery and behavioral strategies could be recruited and maintained for the DMC task, despite extensive retraining. While many LIP neurons show delay period encoding of category during the DMC task, we did not see direction tuning in the average firing rates during the delay period in the DMC task (Fig. 1C, Fig. 5). It is possible that direction is maintained in working memory in LIP populations during the delay period by sparse bursting activity, but not by persistent firing, which cannot be seen by our analysis using single-neuron recordings (Miller et al., 2018). Additionally, previous theoretical work from our lab has demonstrated that recurrent neural networks performing the DMS task may recruit activity-silent computations to compare sample and test stimuli through short-term synaptic plasticity (Masse et al., 2019). In that study using recurrent neural networks trained on both DMS and DMC tasks, delay-period sustained activity was observed more often in tasks which required more complex manipulation of the sample stimulus information compared to the DMS. Our dynamic spike history analysis revealed single-trial dynamics with low-beta frequency oscillatory structure during the delay period in the DMC-only monkeys, but not the pretrained monkeys, which could reflect different working memory dynamics in the DMC-only pair (Miller et al., 2018). Together, this raises the possibility that computations learned during the DMS task which recruited activity-silent working memory during the delay period could explain the observed reduced separability of the sample and test stimulus in the neural subspace in the pretrained monkeys compared to the DMC-only monkeys (Orhan & Ma, 2019).

We also extended the GLM framework to perform dimensionality reduction on neural populations using a flexible tensor-regression model. In complex decision-making tasks, the trials may not be aligned such that common dimensionality reduction methods can be applied without artificially re-aligning single-trial firing rates by stretching or time-warping (Kobak et al., 2016). For example, the touch-bar release ended the trial early at a time determined by the animal in the DMS and DMC tasks. We applied the GMLM to perform dimensionality reduction to find task-relevant features in spike trains when the events in the task were not exactly aligned on every trial, without the need for aligned trial structure. Our approach is related to reduced-rank regression (Steinmetz et al., 2019; Stringer et al., 2019) and the recently proposed model-based targeted dimensionality reduction (Aoi et al., 2020) with two important distinctions: (1) our model is fit to spike trains through an autoregressive Poisson observation model and (2) we consider a more general tensor decomposition of task-related dynamics. The tensor decomposition is used to describe low-rank temporal dynamics in response to stimulus events, similar to low-rank receptive field models of early visual neurons (Ahrens et al., 2008; Park & Pillow, 2013; Elsayed et al., 2020), and those components are shared across all neurons in a population. Unlike PSTH-based dimensionality reduction methods, the Poisson spike generation mechanism accounts for discrete spiking observations and aspects of single-trial dynamics through the spike history kernel and tensor-based dynamic spike history (Park et al., 2014; Holbrook et al., 2017). In contrast to demixed principal components analysis (Kobak et al., 2016) which requires balanced conditions across all variables to recover task-relevant subspaces, the cosine-tuned GMLM takes into account cosine-like direction tuning observed in sensory regions in order to disentangle category and direction information even though direction and category are not separated in the task. Bayesian inference in this model allowed us to quantify uncertainty in the low-dimensional subspace and test hypotheses about the geometry of neural representations. This modeling framework could extend to many other tasks and questions, given appropriate linearizations of specific tasks. For instance, the tensor could be extended to model slow trial-to-trial changes in stimulus response within a recording session by including coefficients for weighting each trial, thereby generalizing applications of tensor component analysis as in Williams et al. (2018).

There are several important limitations about the inferred behavioral strategy and neural mechanisms in the present study. Primarily, this study included only a small number of animals, as is the norm in non-human primate experiments. Furthermore, multiple cortical and subcortical areas are involved in decision making, and our analysis only considered neural activity in LIP. Even within a single region, it is possible that our results could depend on differences in sampling within LIP between animals, or other factors not directly related to the animals’ training history. LIP recording sessions were performed using different sets of motion directions for the two pairs (six directions for the DMC-only monkeys and eight for the pretrained pair). However, we do not believe this small difference in sample directions contributed to the observed differences in LIP because the monkeys were trained using many more motion directions (Sarma et al., 2016). We cannot exclude the possibility that animals could switch behavioral strategy with additional training such that, for example, both pairs of monkeys would perform similarly. Consequently, the possibility remains that LIP representations of the DMC task could change to match the currently adopted behavioral strategy, rather than purely reflecting training history. We think this is unlikely because the LIP recordings were made after all animals had received extensive training on the DMC task and their behavioral accuracy had appeared to asymptote at a high-level of accuracy (Sarma et al., 2016).

There are multiple ways that the brain could learn to perform the same task. Average results across animals may therefore fail to reflect the neural mechanisms of decision making in individual animals (Golowasch et al., 2002; Rahnev & Denison, 2018). Individual differences are a major focus of human decision-making research and have led to many insights into cognitive functions including working memory (Vogel & Awh, 2008; Luck & Vogel, 2013). Here, we explored between subject differences in the dimensionality and the relationship between direction and category tuning in LIP, and we found differences that correlated with long-term training history. Primates in particular may participate in many experiments and receive extensive training in multiple closely related tasks over the course of years. Experimenters should report and consider animals’ training histories when interpreting such data and when comparing seemingly conflicting results from different labs. In conclusion, the low-dimensional dynamics that posterior parietal cortex enlists to support abstract visual categorization can manifest differently across subjects, and exploring long-term effects of training over more subjects can provide broader perspectives of the diverse neural computations that give rise to decision making skills.

## 4 Methods

### 4.1 Data

All datasets used for this study were previously published in Swaminathan & Freedman (2012) and Sarma et al. (2016).

#### 4.1.1 Tasks

The details of the tasks have been described previously for monkeys B and J (DMC-only monkeys; Swaminathan & Freedman, 2012; Swaminathan et al., 2013) and monkeys D and H (pretrained monkeys; Sarma et al., 2016). For all animals, stimuli were high-contrast, 100 % coherent random dot motion stimuli with a dot velocity 12 °s^−1^. The motion patch was 9.0° diameter, and the frame rate was 75 frames/s. Monkeys were required to keep fixation within a 2°radius of the fixation point during each trial.

For monkeys D and H, there were eight sample stimulus directions for both the DMS and DMC tasks, spaced 45° apart: {22.5°, 67.5°, 112.5°, 157.5°, 202.5°, 247.5°, 292.5° and 337.5°}. The test stimuli for the DMS task were 45°, 60°, 75°, or 0° (match) away from the sample stimulus, giving a total of 24 possible motion directions in the task. Test stimuli for the DMC task were the same as the eight sample stimuli. The stimulus presentations were 667 ms and the delay period was 1013 ms.

The DMC task for monkeys B and J used directions spaced evenly in 60°intervals: {15°, 75°, 135°, 195°, 255° and 315°}. The stimulus presentations were 650 ms and the delay period was 1000 ms.

For all monkeys on the DMC task, the motion directions were split evenly into two categories separated by a constant boundary at 45°and 225°.

The DMC task for monkeys B and J included a set of null-direction trials (Fig. S6A). In these trials, the sample direction was along the category boundary (45°or 225°) and the test direction was either 135°or 315°(one direction from each category, furthest from the boundary). These trials were not used to fit neural models, but examined for behavior in Fig. S6B. The monkey’s response was randomly rewarded at 50 % chance on these trials. We note that monkeys B and J were first trained on a simplified DMS task where the sample and test stimuli were either match or 180° opposite. This version of DMS task therefore did not require fine motion direction discrimination, and all correct sample-test response pairs in this task matched were consistent with the DMC task.

#### 4.1.2 Electrophysiology

Neurons in LIP were recorded using single tungsten microelectrodes. During both the DMS and DMC tasks, the motion stimuli were placed inside an LIP cell’s response field.

In this study, we included only cells with a mean firing rate of at least 2 sp/s, averaged from sample stimulus onset to test offset. We included *N* = 31 cells from 26 sessions for monkey b, and *N* = 29 from 27 sessions for monkey J. For monkey D, *N* = 81 cells from 39 sessions for the DMS task, *N* = 63 cells from 33 sessions for the DMC early period, and *N* = 137 cells from 59 sessions for the DMC late period. For monkey H, *N* = 89 cells from 55 sessions for the DMS task, *N* = 106 cells from 40 sessions for the DMC early period, and *N* = 114 cells from 50 sessions for the DMC late period.

#### 4.1.3 Data used for modeling

For all the modeling and decoding analyses, we included only correct trials. The null-direction trials for DMC task for monkeys B and J were not included for model fitting.

We considered a window in each trial starting from sample stimulus onset until 50 ms after the touch-bar release (if a touch-bar release occurred) or 50 ms after the test motion offset. We discretized the spike trains during each trial into 5 ms bins. We note that on non-match trials the animal was required to hold the touch bar until a second test stimulus (which is always a match) appeared. However, the second test stimulus presentation was never included in our analysis.

We used 10 fold cross-validation to compare the models. The trials from each cell were divided into folds evenly by sample directions. For example, if there were 40 trials recorded with sample motion of 22.5° for one cell, these trials were divided into groups of four to make the folds.

For plotting the PSTHs in Fig. 1C, Fig. 3D, and Fig. S8, we smoothed the average spike rate over trials conditioned on motion direction using a Gaussian kernel with a 30 ms width.

#### 4.1.4 Behavioral performance

For quantifying behavioral performance, we only analyzed behavior during the LIP recording sessions. In the behavioral analyses in Fig. 7A and Fig. S6B, we estimated the fraction correct (or fraction touch-bar releases) independently in each condition using a beta-binomial model. The prior parameters in the model were *α* = 1 and *β* = 1 (for a beta distribution over the prior fraction correct or touch-bar released trials). In this model, the posterior over the fraction correct (or touch-bar released) is a beta distribution. The point estimate of the fraction correct was the posterior mean, and the error bars denote 99 % credible intervals over the posterior.

### 4.2 GLM for single cells (the full-rank model)

In this section, we define the generalized linear point-process model of single cells during the DMS and DMC tasks. This class of model for single neurons in decision-making tasks is defined in general in Park et al. (2014). The GLM defines the distribution of spike count at time *t* as a Poisson random variable with mean rate given as a linear function of external events (here, stimulus and touch-bar release) and previous spiking activity:

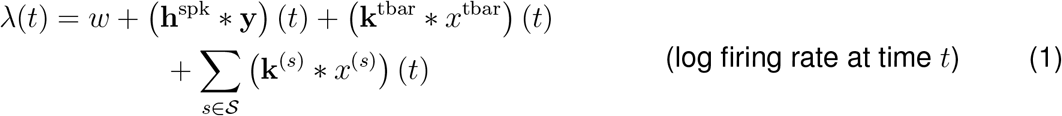

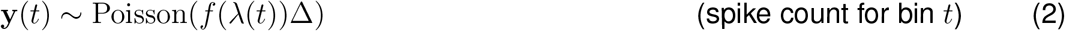

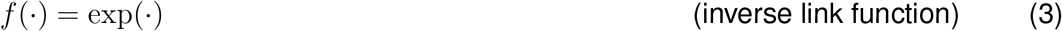

The ∗ operator denotes convolution. The bin width is Δ, and the log baseline firing rate parameter is *w*. Recent spiking activity, **y**, affects the rate through the spike history kernel, **h**^spk^. The stimulus event regressors *x*^(*s*)^(*t*) are functions of at time representing information about the motion stimulus events. The set of all stimulus events is *S*. The touch-bar event is *x*^tbar^. The linear temporal kernels **k**^(*s*)^ and **k**^tbar^ describe the cell’s response to each external variable (stimulus or touch-bar, respectively) as a function of time. The stimulus events we consider encapsulate both sample and test stimuli, but the configuration and number of stimulus kernels depends on the specific model parameterization.

We parameterized the temporal kernels using raised cosine basis functions (Pillow et al., 2008). The stimulus kernel basis consisted of *P*_spk_ = 24 functions with a nonlinear stretching parameter of 0.2 and peaks spanning 0 ms to 1500 ms (Fig. S1J, left). We aligned the basis so that the first basis was zero at exact time of stimulus onset, giving peaks between 40 ms to 1540 ms relative to stimulus onset. The touch-bar basis was constructed using the first *P*_tbar_ = 8 functions of the stimulus basis (Fig. S1J, middle. The functions were reversed and shifted the basis so that the function peaks ranged from −235 ms to 25 ms relative to the touch-bar release and the fastest temporal resolution of the basis set was near the touch-bar release time. We used *P*_stim_ = 24 basis functions for the spike history (Fig. S1J, right). The first two basis functions were Kronecker delta functions to account for the first two bins (0 to 5 ms and 6 to 10 ms after a spike). The remaining eight functions were a raised cosine basis set with nonlinear stretching parameter of 0.05 and peaks from 10 ms to 20 ms post spike.

We define the kernels as the bases times a set of coefficients:

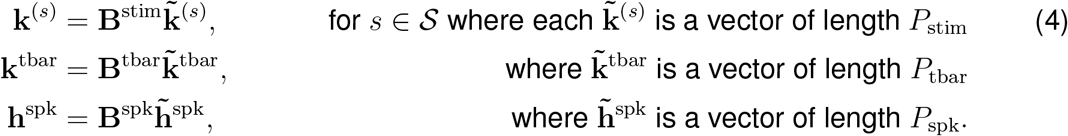

The parameters that are fit to data are 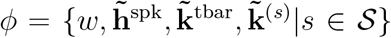. This choice of basis ensures that the stimulus kernels are causal: the stimulus filters only contribute to firing rate after stimulus onset. In contrast, the touch-bar release is acausal: touch-bar release can contribute to the spike rate before the behavior to reflect buildup to the match decision.

We linearized the task events as point events in time. The touch-bar release is given as

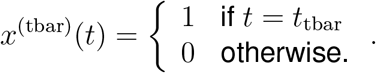

where the time the monkey released the touch-bar to signal a match response is *t*_tbar_ (if the touch-bar was release in the trial). We similarly consider the stimulus onsets (both sample and test) as point events. The sample stimulus duration is constant across all trials, and although the test stimulus is terminated early by a touch-bar release, this does not factor into the window of the trial we model. However, the model can extend to tasks with variable stimulus duration, as has been shown previously (Park et al., 2014). Each GLM kernel gives a scalar contribution to firing rate of the relative time of the task event

We considered a set of four nested models of increasing complexity for the motion stimuli. For simplicity of notation, we present the linearization for a single trial. The sample stimulus onset time is *t*_sample on_ and the test stimulus onset is *t*_test on_. The sample and test stimulus directions are *θ*_sample_ and *θ*_test_ for sample, test ∈ {1, 2, …, *D*} where *D* is the total number of stimulus directions in the task. The stimulus directions belong to categories denoted *c*_sample_, *c*_test_ ∈ {1, 2}.

1. **No category or direction model** (Fig. S1A). This model includes only two stimulus regressors/kernels: one for the sample stimulus onset and one for the test stimulus onset (Fig. S1A): 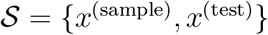. The regressors are defined as point events

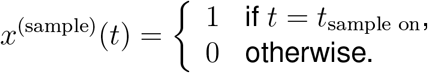

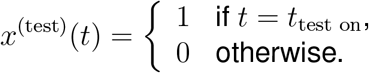

This model captures temporal dynamics in the mean response for each neuron across all stimuli.
2. **Category only model** (Fig. S1B). This model includes four stimulus kernels: two for each category for the sample stimulus (*x*^(cs1)^ and *x*^(cs2)^), and two separate kernels for the test stimulus categories (*x*^(ct1)^ and *x*^(ct2)^). The regressors are again point events, but the points are now conditioned on stimulus category (but not specific direction). For each category *k* ∈ {1, 2}:

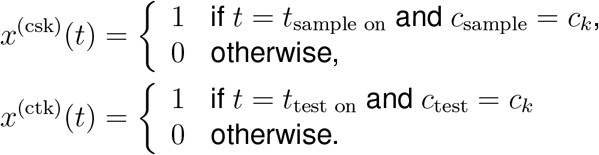

Although the DMS task does not include category, we still applied this model to those data as if there was a category boundary at 45°and 225°.
3. **Cosine direction tuning model** (Fig. S1C). The cosine-tuned model includes both stimulus category and direction tuning, but direction encoding is constrained to a parametric form with cosine tuning. The model includes six stimulus events: two for each category for the sample stimulus (*x*^(cs1)^ and *x*^(cs2)^), two for the test stimulus categories (*x*^(ct1)^ and *x*^(ct2)^), and two for the sine and cosine of the direction (*x*^(sin)^ and *x*^(sin)^). The sample and test category events are defined as in the previous model. The direction regressors are weighted point events, which are shared for both sample and test stimuli:

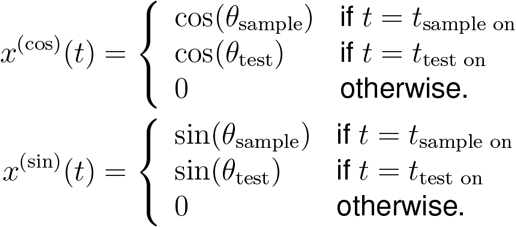
4. **Full model** (Fig. S1D). The full model allows for general (non-cosine) direction tuning. However, we constrained the model to have the same direction tuning for both sample and test stimuli; that is, the difference in tuning between two directions within the same category was the same for both sample and test stimuli. The model included one stimulus regressor event for each directions plus two test category events (for the DMC task with *D* = 6 stimulus directions, there are eight kernels) For each trial and direction *θ*_*d*_ for *d* ∈ {1, 2, …, *D*}, the stimulus regressors are

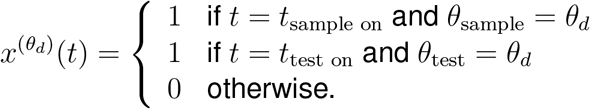

The two stimulus events that parameterize the test category responses (*x*^(ct1)^, and *x*^(ct2)^) are defined as before. This parameterization maintains identifiability: it would not be identifiable to directly expand the cosine model to have a kernel for each direction plus two sample and two test category kernels. As a result of the identifiability constraint, the interpretation of the corresponding kernels is different compared to the cosine tuning model: in this model, 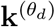 is the kernel for a stimulus in the *θ*_*d*_ direction plus the response to a stimulus of the category of *c*_*d*_. Therefore, we view **k**^(ct1)^ as the kernel to a test stimuli of category 1 minus the kernel for a category 1 sample stimulus (thereby subtracting the sample category tuning away from the direction kernel and adding the test category tuning).

We considered two additional models included in supplementary analyses that included independent sample and test direction tuning (Fig. S6). The independent direction cosine tuning model had eight kernels total: four for the sample and test category, four for the cosine and sine weights of the sample and test directions. The full independent direction model simply had one kernel for each sample direction and one kernel for each test direction These two models are defined analogously to the common direction tuning models.

#### 4.2.1 Prior probabilities over model parameters

We defined zero-mean Gaussian priors over the stimulus kernels. The orthonormal basis functions controlled temporal smoothness of the kernels, and the prior distributions were independent over time. We defined i.i.d. priors for the *i*th coefficients of the stimulus kernels (i.e., a prior over the set of 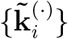 for each *i* ∈ {1, …, *P*_spk_}). We describe the priors of the stimulus kernels for each of the nested models.

1. **No category or direction model**. We considered that the sample and test kernels would likely be correlated if they reflect the dynamics of common bottom-up sensory input. To construct a correlated prior, we assumed that the kernel could be constructed as the sum of stimulus-independent kernel (a response purely to contrast or motion in general), and a sample kernel or a test kernel (responses to the task epoch):

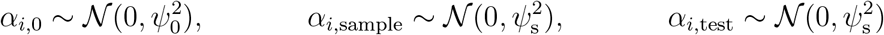

such that

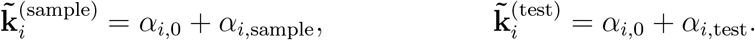

Using the rules of linear transformations of Gaussian variables, we obtained a correlated Gaussian prior over the original two sample and test kernels. The set of hyperparameters was 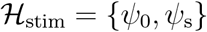.
2. **Category only model**. For the category-dependent kernels, we made a similar Gaussian construction

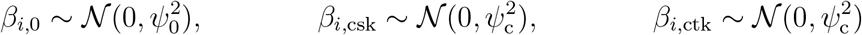

such that

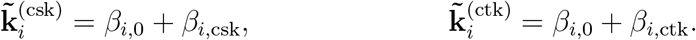

The set of hyperparameters was 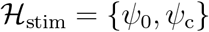.
3. **Cosine direction tuning model**. We used the same priors for the four category kernels as in the category-only model. We placed an independent Gaussian prior over the cosine and sine weights:

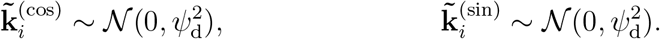

The set of hyperparameters was 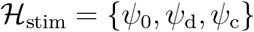.
4. **Full model**. For constructing the prior over individual direction kernels, we assumed that direction tuning should be smooth as a function of angle. We therefore used a Gaussian process prior over the direction weights. To nest the cosine-tuning model in the full model and provide regularization, we also included latent sine and cosine direction weighting. Sample category weights were included as before. We define the pieces of the prior as

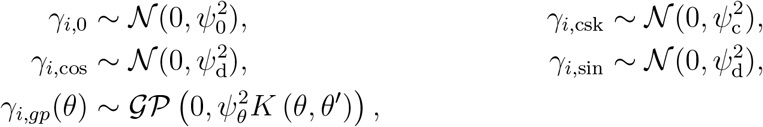

where the Gaussian process kernel over angle is (Padonou & Roustant, 2016)

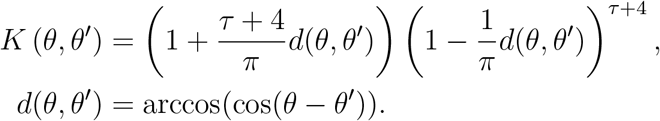

The hyperparameter *τ* ≥ 0 determined the arc length over which similar directions are correlated, similar to a length scale in Gaussian process kernels on the real line. The complete direction plus sample category kernel was then defined as for each direction *d* ∈ {1, …, *D*}

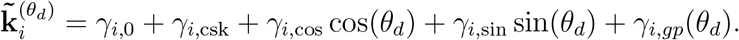

The test category prior was defined slightly differently than in the previous two models due to the identifiability constraints on our parameterization. We defined the prior using the construction

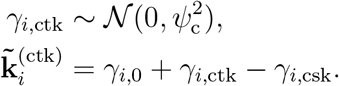

Because 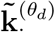 and 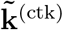 were again simply linear functions of Gaussian variables, we obtained a Gaussian prior with zero mean for the kernels depending on the hyperparameters set 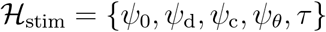

For the supplementary models with independent sample and test direction tuning, the priors followed the same construction as above. The direction hyperparameters were shared for the sample and test direction kernels.

We placed an i.i.d. Gaussian prior on the spike history and touch-bar coefficients

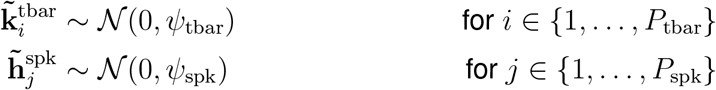

The prior over *w* was the improper uniform prior: *p*(*w*) ∝ 1.

The complete set of hyperparameters was therefore 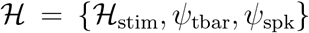. We defined hyperpriors over each hyperparameter independently as half-*t* distributions (Gelman, 2006). For each 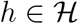

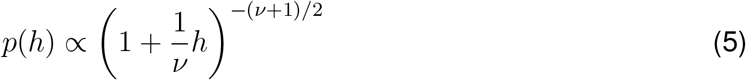

where we set *ν* = 4.

#### 4.2.2 MAP estimation with evidence optimization

We fit the GLMs to each LIP cell using MAP estimation. To set the hyperparameters for the GLM, we used an approximate evidence optimization procedure (Sahani & Linden, 2003; Park & Pillow, 2013; Zoltowski & Pillow, 2018). We used a Laplace approximation of the posterior over model parameters to get likelihood of data given hyperparameters to estimate the log evidence (i.e., the marginal distribution of the data given the hyperparameters). We then optimized the log posterior over the hyperparameters. Because the hyperparameters are constrained to be positive, we optimized the log-transformed hyperparameters. We set the hyperparameters and parameters of the GLM independently for each fold of cross-validation.

Specifically, we maximized an approximation of the log posterior of the hyperparameters given the data. The posterior is

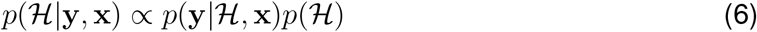

and we want to find

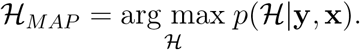

The evidence term can be written using Bayes’ rule as

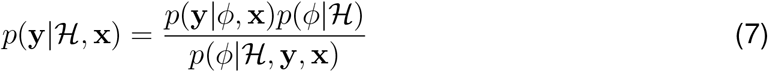

where *ϕ* denotes the model parameters. The posterior over the parameters 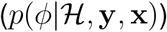 is only given up to an intractable normalizing constant. We therefore took a Laplace approximation of the posterior distribution over parameters. The Laplace approximation was a Gaussian distribution centered around the MAP estimate of the parameters

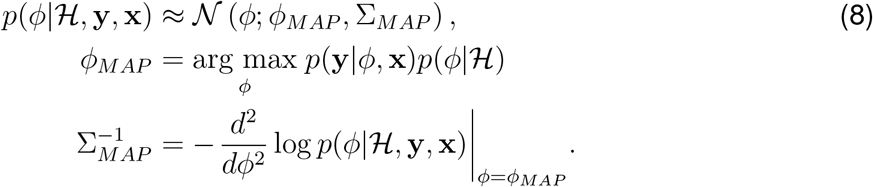

The MAP estimate given the hyperparameters, *ϕ*_*MAP*_, was found numerically (the log posterior over the parameters is log concave). Given this approximation, we evaluated the right side of Eq. 7 at *ϕ*_*MAP*_. We then maximized the log posterior over the hyperparameters (Eq. 6) numerically to find 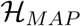. The final MAP estimate of the models parameters was *ϕ*_*MAP*_ given 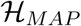.

### 4.3 GMLM definition

The GMLM is a special case of the GLM in which the linear kernels in a population of neurons are assumed to share low-dimensional structure, rather than being modeled independently. In general, the model is a GLM in which the regressors and parameters (or a subset thereof) from all the neurons in a population can be expressed as tensors (or multi-way arrays). The parameters (or a subset of the parameters) are then assumed to have a low-rank structure: the parameter tensor can be decomposed into a small number of components. We emphasize that the neurons need not be simultaneously recorded for this model: we can still fit the parameters if only one neuron is observed at each time point.

Here, we define the GMLM for the DMC task. Introducing an index for neuron *n* ∈ {1, 2, …, *N*}, we defined the model for the spike count in bin *t* ∈ {1, 2, …, *T*} for neuron *n* as

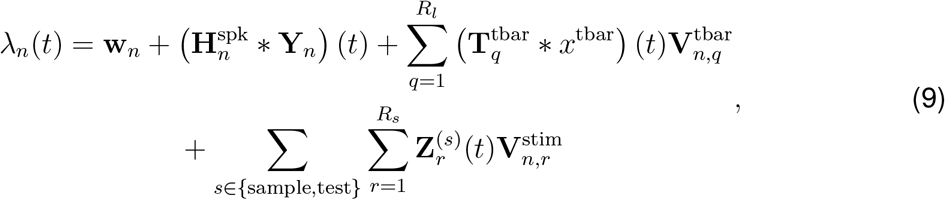

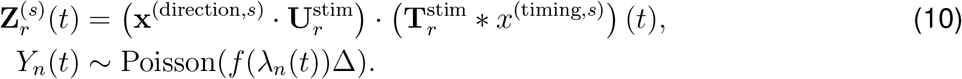

The matrices 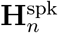, **T**^tbar^, and **T**^stim^ denote matrices whose columns are temporal kernels for the stimulus, touch-bar and spike history respectively. Subscripts of those matrices indicate a particular column or kernel. Similarly, **H**^spk^ contains the spike history kernels for each cell. The length *N* vector **w** contains the baseline firing rates for each neuron. The baseline firing rates and spike history kernels are equivalent to the single-cell GLM.

We note that in this model, both the regressors and the parameters are decomposed into components (for the stimulus parameters **T**^stim^, **U**^stim^, and **V**^stim^ and regressors **x**^(direction,*s*)^ and *x*^(timing,*s*)^), and thus we have a simple multilinear form for the stimulus tuning rather than writing out a dense tensor.

The touch-bar kernels are parameterized as low-rank matrix factorization where **T**^tbar^ contains *R*_*l*_ temporal kernels and **V**^tbar^ is a matrix of neuron loading weights of size *N* × *R*_*l*_. In our notation, 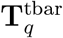 denotes the *q*th column or kernel, and 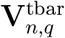 is the element in the *n*th row, *q*th column of **V**^tbar^. The touch-bar subspace is the span of the columns of **V**^tbar^. Thus, the model effectively approximates the GLM touch-bar kernel for neuron *n* as 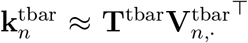. The touch-bar function, *x*^tbar^, is the same as in the GLM.

The stimulus kernels were parameterized as a tensor factorization of rank *R*_*s*_. As we did for the GLM, we defined a matching set of nested models to parameterize the DMC task. As with the GLM definitions, we defined the regressors for a single trial for simplicity of notation. The notation for the stimulus timing and directions are the same as in the GLM definition.

The set of temporal kernels, **T**^stim^, did not depend on the stimulus direction or category. The temporal regressors were point events representing the stimulus onset time for each *s* ∈ {sample, test}. These were the same for all GMLM parameterizations (Fig. S1E):

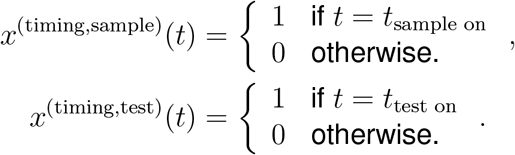

The set of stimulus weights, **U**^stim^, was a matrix *S* × *R*_*s*_ of coefficients for the particular stimulus identity (for example, weights to encode sample, test, direction, and category) where *S* is the same as the number of stimulus kernels in the matching GLM. Each observation had two stimulus direction regressor vectors: **x**^(direction,sample)^ and **x**^(direction,test)^. The entries of the stimulus direction regressors (**x**^(direction,*s*)^) mirrored the kernel structure in the GLM parameterizations (this vector is constant for all *t* in a single trial). The stimulus direction coefficients depended on the model.

1. **No category or direction model** (Fig. S1F This model contained two stimulus regressor elements indexed by {sample, test}, As with the GLM, these elements represent the identity of a stimulus event as sample or test, but does not include category or direction information.

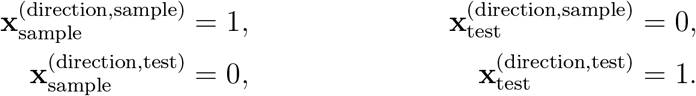
2. **Category only model** (Fig. S1G) This model includes four stimulus direction regressors representing the stimulus category and whether it is sample or test. For the indices {cs1, cs2, ct1, ct2}, the regressors are

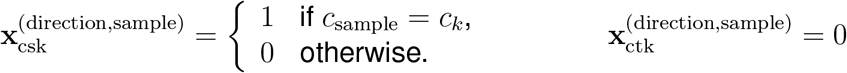

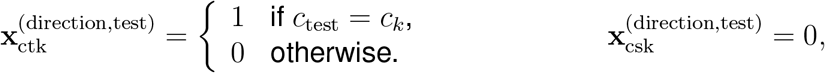

for category *k* ∈ {1, 2}.
3. **Cosine direction tuning model** (Fig. S1H). The cosine-tuning model included six stimulus regressors representing the identity of a stimulus event as sample or test, the motion category, and the sine and cosine of the direction. The regressors are indexed by {cs1, cs2, ct1, ct2, cos, sin}. The category terms are the same as in the above model. The direction tuning components are defined as cosine and sine weights of the direction:

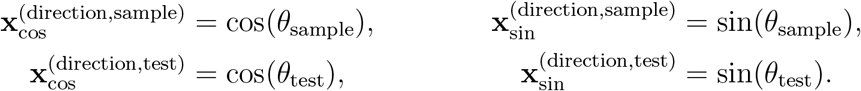
4. **Full model** (Fig. S1I). This model includes one regressor for each stimulus direction and two for the test stimulus category indexed by the *D*+2 coefficients in {ct1, ct2, *θ*_1_, …, *θ*_*D*_}. For each *d* ∈ {1, …, *D*}

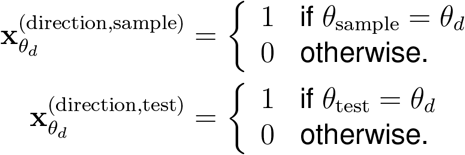

The test category regressors (indexed by ct1 and ct2) are the same as in the previous two models. As with the full GLM, the specific model construction does not include additional weights for the sample category for identifiability. The coefficients in **U**^stim^(*θ*_*i*_) represent the tuning strength for direction *θ*_*i*_ plus the tuning for the category of *θ*_*i*_. Therefore, the coefficients in **U**^stim^(ctk) represent the tuning for a test stimulus of category *k* minus the tuning for sample stimulus of category *k*.

Together, the matrices **T**^stim^, **U**^stim^, and **V**^stim^ define a CP or PARAFAC decomposition of the GLM stimulus kernels over a population of cells (Kolda & Bader, 2009). That is, the *s*th stimulus kernel at time *t* for neuron *n* is approximated as the low-rank decomposition

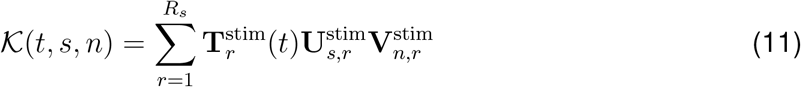

Another way to view the dimensionality reduction is that the values of **Z**.^(*s*)^(*t*) give an *R*_*s*_ dimensional representation of the response to each stimulus over time. Each neuron’s response to the stimulus is given as a linear projection of that low-dimensional stimulus with weights defined as the rows of the matrix **V**^stim^ so that **V**^stim^ defines the stimulus subspace.

The temporal kernels of the GMLM are parameterized using the same basis set as the GLM:

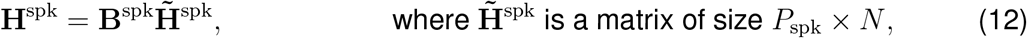

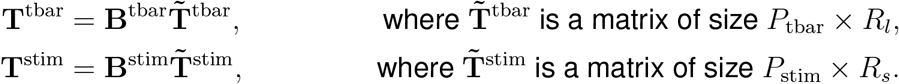

We set *R*_*l*_ = 3 for all GMLM fits and we selected *R*_*s*_ using cross-validation (see Section 4.3.3). The set of parameters that are fit to the data from all the trials in an LIP population is 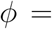 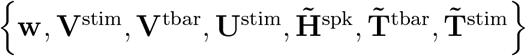.

For Bayesian inference, we defined prior distributions the same way we did for the GLM. The prior for the stimulus kernels was defined independently for each component *r* of the stimulus direction regressor matrix (i.e., each column 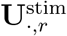). The vectors 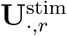 and corresponding GLM kernel parameters 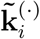 are the same length and are indexed by the same set of stimulus events The prior over the vector 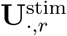 was therefore the same as the prior over the 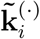 for the corresponding GLM. The same stimulus hyperparameter set 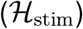 was used for each model parameterization. However, unlike the single-cell GLM fits, the hyperparameters were shared across all neurons in each LIP population.

The prior for the entries of the spike history kernels, 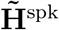, was i.i.d. normal with zero mean and variance 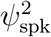. Similarly, the prior for the entries of the touch-bar kernels, 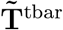, was i.i.d. normal with zero mean and variance 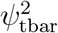. The prior distribution on the entries of neuron loading matrices (**V**^stim^ and **V**^tbar^) and **T**^stim^ was i.i.d. standard normal. We again used an improper uniform prior on **w**.

The complete hyperparameters set was 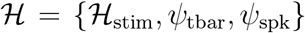. The hyperpriors were the same half-*t* distributions used for the GLM (Eq. 5). We note that these priors only affected the GMLM in the MCMC analysis, as rank selection used the maximum likelihood estimate.

#### 4.3.1 Dynamic spike history

We augmented the log rate in the GMLM with low-rank dynamic spike history components to allow spike history to change over time relative to task events:

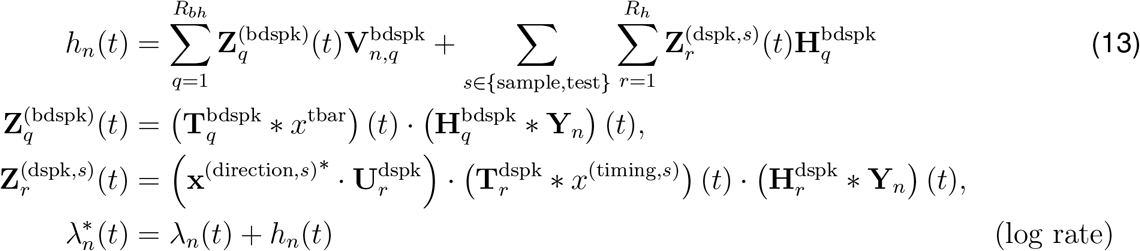

where *λ*_*n*_(*t*) is given by Eq. 9. For completeness, we included two dynamic spike history tensors to mirror the mean-rate filter terms: one for the motion stimuli and a second for the touch-bar release. However, we found including the touch-bar filters provided little improvement to the model’s performance.

The dynamic spike history kernels **H**^dspk^ for the stimulus-dependent spike history (or **H**^bdspk^ for the touch-bar kernel) are shared across all neurons in a population. The stimulus kernels **T**^dspk^ (or **T**^bdspk^ for the touch-bar release kernel) control the contribution of the dynamic spike history kernel relative to stimulus onset (or touch-bar release). As with the stimulus filter tensor, we allow the dynamic spike history kernels to depend on stimulus information through the stimulus scaling terms **U**^dspk^. For simplicity, we limited the stimulus scaling for the dynamic spike history in 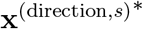 to include only sample or test information as defined for the “No category or direction model” in the previous section. We found that including category or direction information did not significantly affect our results (results not shown). Each neuron weights the dynamic spike history components by the loading matrices **V**^dspk^ and **V**^bdspk^.

At any given time in the trial, the spike history for a neuron is still a linear function of past spiking. The “effective” spike history kernel for a neuron *n* at time *t* can be computed by rearranging the terms in Eq. 13 and adding the constant spike history kernel:

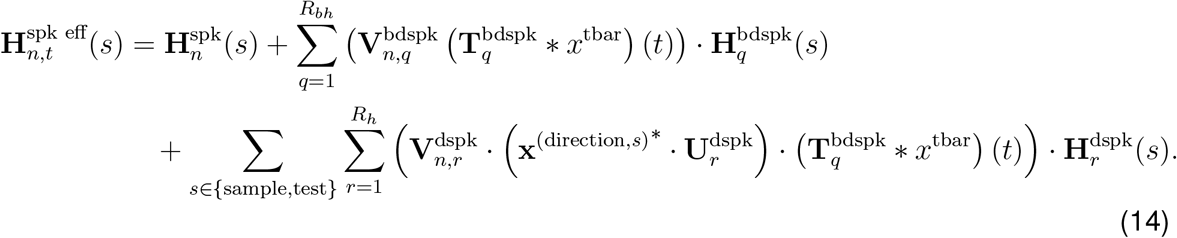

The temporal kernels were parameterized using the same basis set as before:

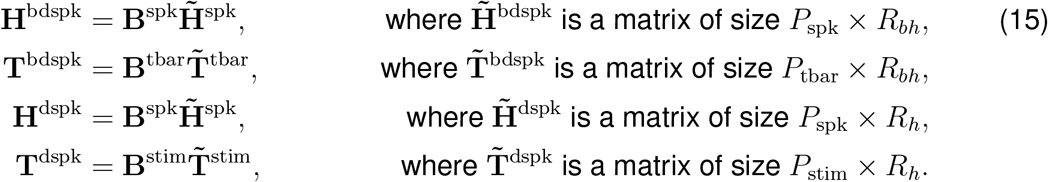

The parameter set for the dynamic spike history model was 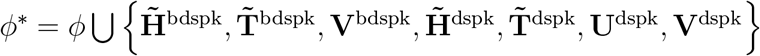.

We set i.i.d. standard normal priors for **V**^dspk^, **V**^bdspk^, 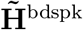, 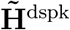 and 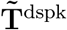. The prior for 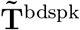 was i.i.d. zero-mean normal with variance 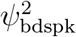. The Gaussian prior for **U**^dspk^ was defined analogously to the stimulus term (for the no category model) with hyperparameters 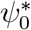 and 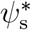. The complete hyperprior set was then 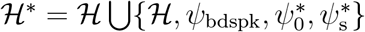.

For all dynamic spike history models here, we set *R*_*bh*_ = 1 and we varied *R*_*h*_ from 1 to 2. For the stimulus filter tensor, we used the cosine direction tuning model with the rank selected in Fig. 4.

#### 4.3.2 Model inference

We performed rank selection in the GMLM by testing cross-validated model performance of the maximum likelihood fit. We used gradient-descent methods to numerically maximize the log likelihood for fold. We initialized the GMLM components randomly. The entries of **V**^stim^, **V**^tbar^, and 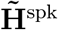 were generated as standard normal. The matrices 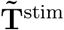 and 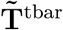 were random orthonormal matrices. The baseline firing rate parameters, *w*, were drawn independently from a normal distribution.

For the MAP estimates shown in Fig. 3 and Fig. 5, we set the hyperparameters to the marginal posterior medians of each hyperparameter estimated using Markov chain Monte Carlo methods (described below). We then maximized the posterior log likelihood given those hyperparameters.

For the Bayesian analyses of the GMLM, we used MCMC to generate samples from the posterior distribution of the model parameters and hyperparameters given all data from an LIP population. We used Hamiltonian Monte Carlo (HMC) to sample jointly from the posterior of the parameters and the log-transformed hyperparameters. The log transform on the hyperparameters ensures that the hyperparameters are positive. A detailed description of the HMC sampling algorithm is given in (Neal, 2011). The Hamiltonian equations were solved numerically using a leap-frog integrator with step size *ϵ* for *S* steps. We set 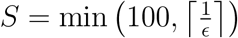 where ⌈ · ⌉ denotes the ceiling operator. The maximum number of steps was 100 to limit computational costs per sample. However, after tuning the sampler during warmup, we found that *S* < 100.

We denote the vectorized set of all parameters and log-transformed hyperparameters for sample *s* as 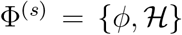. We initialize the model parameters for the sampler (*s* = 1) by initializing the parameters randomly as we did for the maximum likelihood estimation. The log hyperparameters were initialized as i.i.d. draws from a standard normal distribution. The HMC sampler requires specifying a *P* × *P* mass matrix, **M**. Because the model is high dimensional, we assume **M** is diagonal.

We tuned the parameters of the sampler (*ϵ* and **M**) by generating 25 000 warmup samples (also known as “burn-in”). The initial value of the step size was *ϵ* = 0.01. We used the dual-averaging algorithm of Nesterov (2009) to adapt *ϵ* at each step for the first 24 000 warmup samples (and fixed for the last 1000 warmup samples) to achieve a desired acceptance rate. We used the parameters given in Hoffman & Gelman (2014) to control the learning rate and target sample acceptance rate (80 %). The mass matrix **M** was set at three steps.

1. Sample 1: **M** is initialized as identity matrix.
2. Sample 4001: 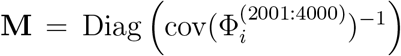. The diagonal is the inverse empirical variance of each parameter given samples 2001 to 4000.
3. Sample 19 001: 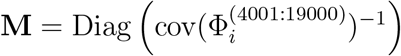.

After warmup, we generated 50 000 HMC samples. These sampled were used as the estimate of the posterior distribution of the model parameters and hyperparameters.

One source of autocorrelation in the HMC sampler that could reduce the quality of inference is that the GMLM tensor components could be re-scaled without changing the likelihood. For any *a, b* ≠ 0, the *r*th component of the GMLM stimulus kernel tensor can be rescaled

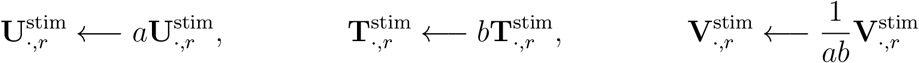

without changing the resulting kernel tensor. Thus, the log likelihood remains constant. One way to is to constrain fix the norm of two of those vectors, and thereby disallowing re-scaling. Inference can then be performed for those parameters on an appropriate manifold (product of sphere manifolds) using geodesic Monte Carlo methods (Byrne & Girolami, 2013; Holbrook et al., 2016). Instead, we took a different approach by including an efficient MetropolisHastings (MH) step for rapidly traversing the locally flat region of the likelihood without additional constraints on the model parameters. The MH step was performed independently for each component *r*. We define for the current sample *s*

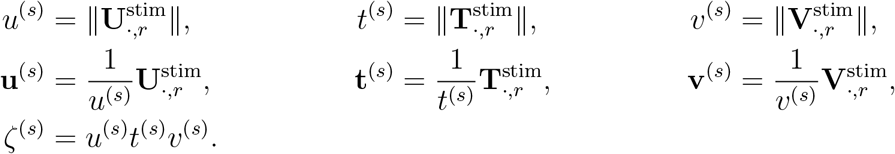

The prior probabilities for each 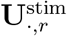, 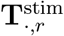, and 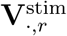 are multivariate Gaussian with zero mean. Therefore, the prior probability of the vector lengths *p*(*u*^(*s*)^, *t*^(*s*)^, *v*^(*s*)^ |**u**^(*s*)^, **t**^(*s*)^, **v**^(*s*)^, 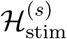) can be factorized into independent chi distributions:

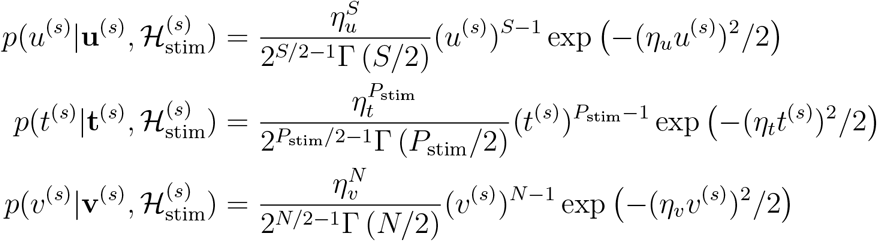

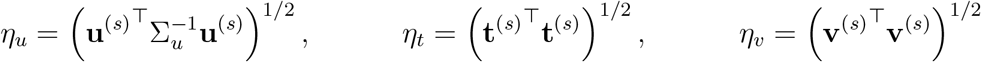

where Σ_*u*_ is the prior covariance matrix for 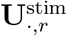 given the hyperparameters 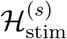 (the Gaussian priors for the other two vectors have identity covariance). Our goal is to construct a MH proposal to focus on the case where the total component norm *ζ*^(*s*)^ is constant. Therefore, we perform a change of variables on the prior over *p*(*u*^(*s*)^, *t*^(*s*)^, *v*^(*s*)^|**u**^(*s*)^, **t**^(*s*)^, **v**^(*s*)^, 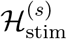) to *p*(*u*^(*s*)^, *t*^(*s*)^, *ζ*^(*s*)^|**u**^(*s*)^, **t**^(*s*)^, **v**^(*s*)^, 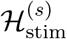) in order to compute

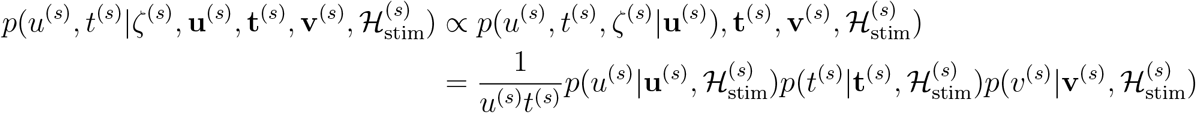

We can then generate independent scaling factors to perform a random walk on the scaling factors:

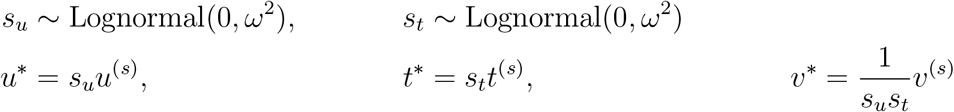

We then accept the proposal *u*\*, *t*\*, *v*\* with the MH acceptance probability

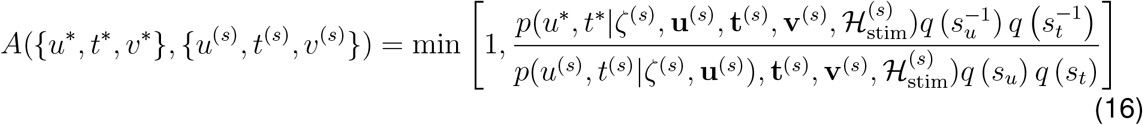

where *q*(*s*) = Lognormal(*s*; 0, *ω*^2^). Because the likelihood remains constant in this proposal, we only need to compute the prior to determine the MH acceptance probability. As a result, this step is very fast to compute. We applied the same class of MH proposal to the touch-bar components. We set *ω* = 0.2 and interleaved 10 MH steps for each tensor component between every HMC step.

#### 4.3.3 Rank selection

We applied cross-validation to select the stimulus kernel tensor rank (*R*_*s*_) for the GMLM. To do so, we computed the mean test log likelihood per trial per cell. For neuron *n*,

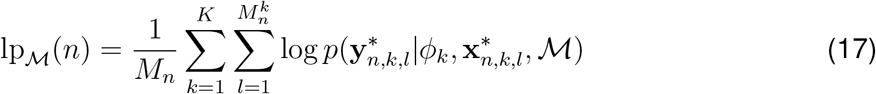

where *K* is the number of folds (*K* = 10 for all the analyses conducted here). The trials in the test set are given as 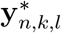 and 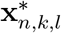 which represent the spike train and regressors respectively for test trial *l* in fold *k*. The number of test trials in fold *k* for the neuron is 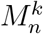, and the total number of trials is 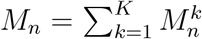. The model parameters for the model 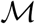 fit to the training data for fold *k* is *ϕ*_*k*_.

We then took the average across all cells

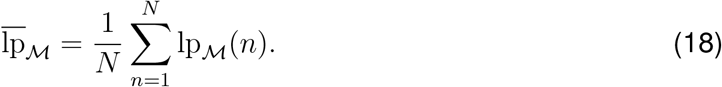

For normalization, we subtracted the 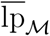 of the GMLM without any stimulus terms (the “rank 0” model):

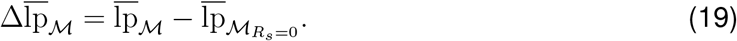

Fig. 4A shows the 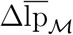 for each GLM and GMLM of from *r* = 1 to 12.

The fraction of log likelihood explained by the GMLM was computed relative to the full GLM (the “full rank” model, denoted 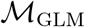):

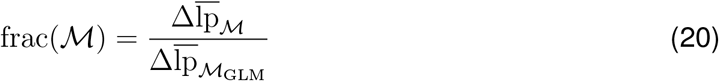

Fig. 4B shows the frac 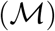 for each GLM and GMLM of from *r* = 1 to 12 for the monkey D, DMC late population. We selected the rank *r* for the full GMLM by selecting the smallest *r* for which frac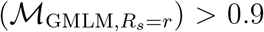 (i.e., the number of model components needed to explain 90 % of the likelihood that could be explained by this GLM framework).

The error bars over the cross-validated log likelihood in Fig. 4A were computed by computing 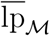 on the test trials for each fold separately (instead of averaging over all *K* folds). The error bars show two standard errors of 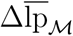 over the folds.

For the dynamic spike history, we compared model predictive performance with leave-one-out cross-validation estimated with Pareto-smoothed importance sampling using the MCMC samples (Vehtari et al., 2017). The leave-one-out cross-validated log likelihoods were computed for each trial, and then we computed the mean cross-validated log likelihood for each neuron.

#### 4.3. Visualizing the GMLM parameters

Fig. 3A shows the individual components of the MAP fit of the full GMLM, which included kernels for each stimulus direction and two kernels for the test stimulus category. For scale, we normalized each component by placing the magnitude of each tensor component in the neuron loading dimension:

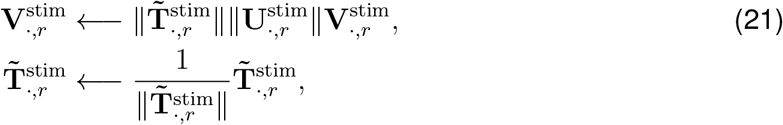

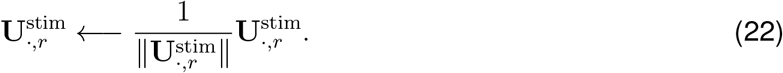

The *r*th row of the left column of Fig. 3A shows the re-scaled 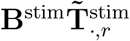. The middle column shows those temporal kernels scaled by the direction weights: the *r*th row plots 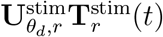 for each direction *d*. The right columns shows the temporal kernels scaled by the additional category weights for the test stimulus: thr *r*th row plots for 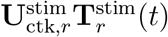 for both categories *k*. The loading weights in the box plot of Fig. 3B show the elements 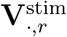 for each component *r*.

The sample stimulus kernels for the example cells in Fig. 3C are the sample direction kernels scaled by the neuron’s loading weights for each component. The *r* row for example neuron *n* shows 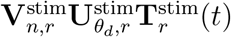 for each direction *d*. The total GMLM tuning (Fig. 3D top row) was the sum over the *r* components.

To visualize the subspaces, we projected the components of the full GMLM into the top three dimensions. The loading weights of the tensor decomposition used to define the model (**V**^stim^) are not constrained to be orthonormal (as is standard for the PARAFAC decomposition). Therefore, we applied a Tucker decomposition (i.e., higher-order singular value decomposition) to find the three-dimensional subspace that captures most of the population’s stimulus tuning structure. We took the stimulus kernel tensor of 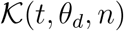 (Eq. 11) for all sample stimulus directions. We then took the Tucker decomposition of the stimulus kernel tensor such that

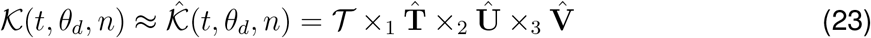

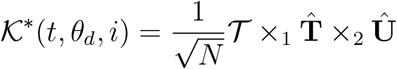

where 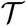 is the core tensor of size *R*_*s*_ × *D*_sample_ × 3, and 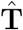, 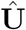, and 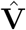 are orthonormal matrices. The filter tensor projected into the top three subspace dimensions (*i* ∈ {1, 2, 3}) for each direction over time is then 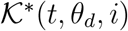.

To find the mean-removed space, we took

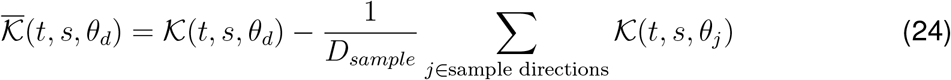

we performed the Tucker decomposition on 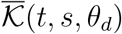 to obtain the mean-removed subspace.

For visualizing the rank-1 dynamic spike history components in Fig. 8B, we plot the posterior median and pointwise 99 % credible intervals computed using MCMC for the normalized temporal filters, **T** ^dspk^*/*|| **T**^dspk^|| and **H** ^dspk^*/*|| **H**^dspk^||. Because t he sign of i ndividual components in the PARAFAC decompositions is not identifiable, we set the sign of the posterior median components with the following transformation in order to better compare across populations:

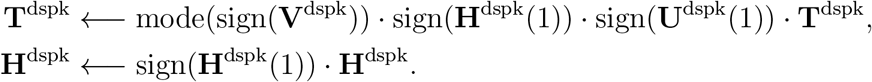

To quantify the timescales of the dynamic spike history kernels (for the pretrained monkeys only), we fit the MAP estimate of the rank-1 dynamic spike history kernel with an exponential function with a least-squares fit.

#### 4.3.5 Bayesian analysis of subspace geometry

We defined tuning metrics in the low-dimensional space estimated by the GMLM with cosine direction tuning to analyze the geometry of task encoding. The metrics were constant over rotations and translations of the latent subspace. We used the posterior distribution of the model parameters estimated using MCMC to establish credible intervals over the metrics.

At each time point *t* relative to stimulus onset, the cosine-tuned GMLM defines direction tuning in the population as an ellipse embedded in 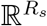 parameterized by angle as

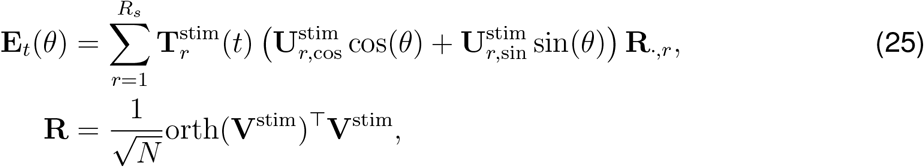

where orth(**V**^stim^) denotes a matrix whose columns contain an orthonormalized basis for the span of the columns of **V**^stim^. Here, the orthogonalized *R*_*s*_-dimensional output space, **R**, is normalized by the number of cells. We computed the angle of the major axis of the ellipse as *θ*_max_ where

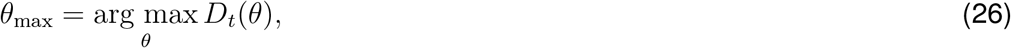

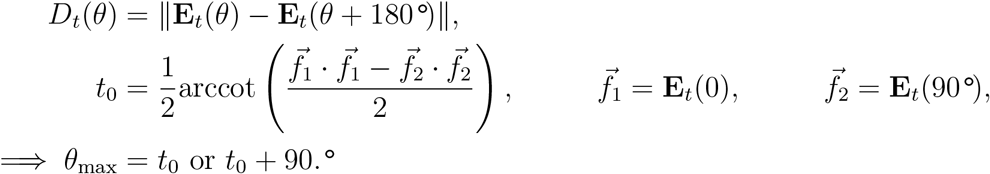

Because *θ*_max_ is identifiable only up to a factor of 180°, we added the constraint *θ* _max_ ∈ [45°, 225°] to relate the angle to category in the task. The norms of the major and minor axes are *D*_*t*_(*θ*_max_) and *D*_*t*_(*θ*_max_ + 90°) respectively.

The category tuning vector is the difference in the low-dimensional tuning space between the category one and category two kernels:

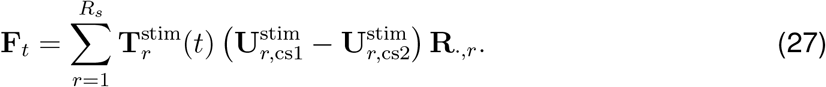

Category tuning norm at each time *t* relative to stimulus onset is then the norm of the vector, ||**F***_t_*||.

For the Bayesian analysis, we computed *θ*_max_, *D*_*t*_(*θ*_max_), *D*_*t*_(*θ*_max_ + 90°), and ||**F**_*t*_|| for each sample from the posterior distribution of the model parameters. We then computed the posterior median and a 99 % credible interval covering 0.5 % to 99.5 % of the posterior for each time *t*.

For the supplementary analyses in Fig. S3 and Fig. S7, we performed component-wise analyses of the GMLM fits. We note that the GMLM posterior has multiple modes: the order of the components can be permuted or a sign flip could occur between **U**^stim^ and **T**^stim^. These modes define equivalent subspaces and kernel tensors, and the prior distributions are the same at each mode. We did not find that the HMC sampler jumped between these modes, and thus we could simply analyze the individual components of the GMLM tensor. For the component-wise analysis, we looked at each *r* ∈ {1, …, *R*_*s*_} individually. The direction tuning for the component was

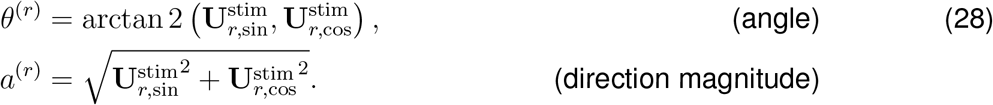

The sample and test category tuning for the component was

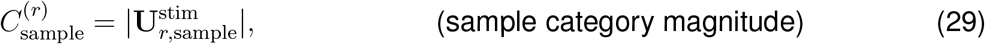

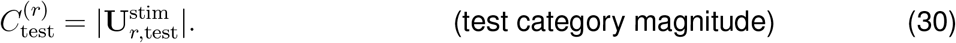

### 4.4 Decoding analyses

All decoders were linear, binary classifiers on pseudopopulation trials spike counts fit with logistic regression in MATLAB using the `fitclinear` function. The training set spike counts were z-scored and the decoder was fit with ridge regression with penalty 0.1. Because the neurons were recorded independently, we constructed pseudopopulation of 50 trials per stimulus. Each pseudopopulation trial consisted of one randomly sampled (with replacement) trial from each neuron in the recorded population for a particular stimulus direction. We repeated the decoding analysis on 1000 random pseudopopulations to obtain bootstrapped confidence intervals.

To decode sample category as a function of time from stimulus onset, we fit and tested decoders using spike counts in a sliding 200 ms window (centered at the decoding time). To test for direction-independent category tuning, training and validation conditions were trials from different directions to test direction-independent category encoding (Sarma et al., 2016). We therefore fit two decoders, each using trials only from a subset of motion directions. Generalization was evaluated using the withheld directions for each decoder, and the total generalization performance was averaged across the two decoders. The two sets of monkeys had a different set of sample directions, and thus different train/validation conditions For monkeys B and J, each training set contained two motion directions, spaced 180° apart: {15° and 195°} and {75°and 225°}. Test sets were then the four remaining motion directions in each condition (135°and 315°trials were in the validation set for both decoders). For monkeys D and H, the training sets were {67.5°, 112.5°, 247.5°and 292.5°} or {157.5°, 202.5°, 337.5°and 22.5°}.

For decoding category decoding during the test stimulus, we used pseudopopulation spike counts in a window from 0 to 200 ms after test motion onset. For these decoders, the training and validation sets included pseudopopulation trials from all motion directions. The decoders were trained using only match (or non-match) trials and tested for generalization on non-match (or match). The total performance was the average across the match-trained and non-match-trained decoders. We trained separate decoders for sample and test category. The DMS populations were excluded from this analysis, because the test stimulus directions depended on the sample stimulus.

### 4.5 Modeling software

All GLM and GMLM analyses were performed using custom software for MATLAB (MathWorks) and CUDA (Nvidia). The GMLM tools are available publicly at https://github.com/latimerk/GMLM_dmc. Tucker decomposition for visualizing the subspaces was performed with Tensor Toolbox for MATLAB (Bader et al., 2019).

## Acknowledgements

This work was supported by a Chicago Fellowship (KWL) and grants NIH R01 EY019041 and DOD VBFF (DJF). We thank Rheza Budiono, Jeffrey Johnston, Pantea Moghimi, Barbara Peysakhovich, Jonathan Pillow, Matthew Rosen, Jacob Yates, and Oliver Zhu for helpful comments and discussions.

## Supplementary Information

**Figure S1:**
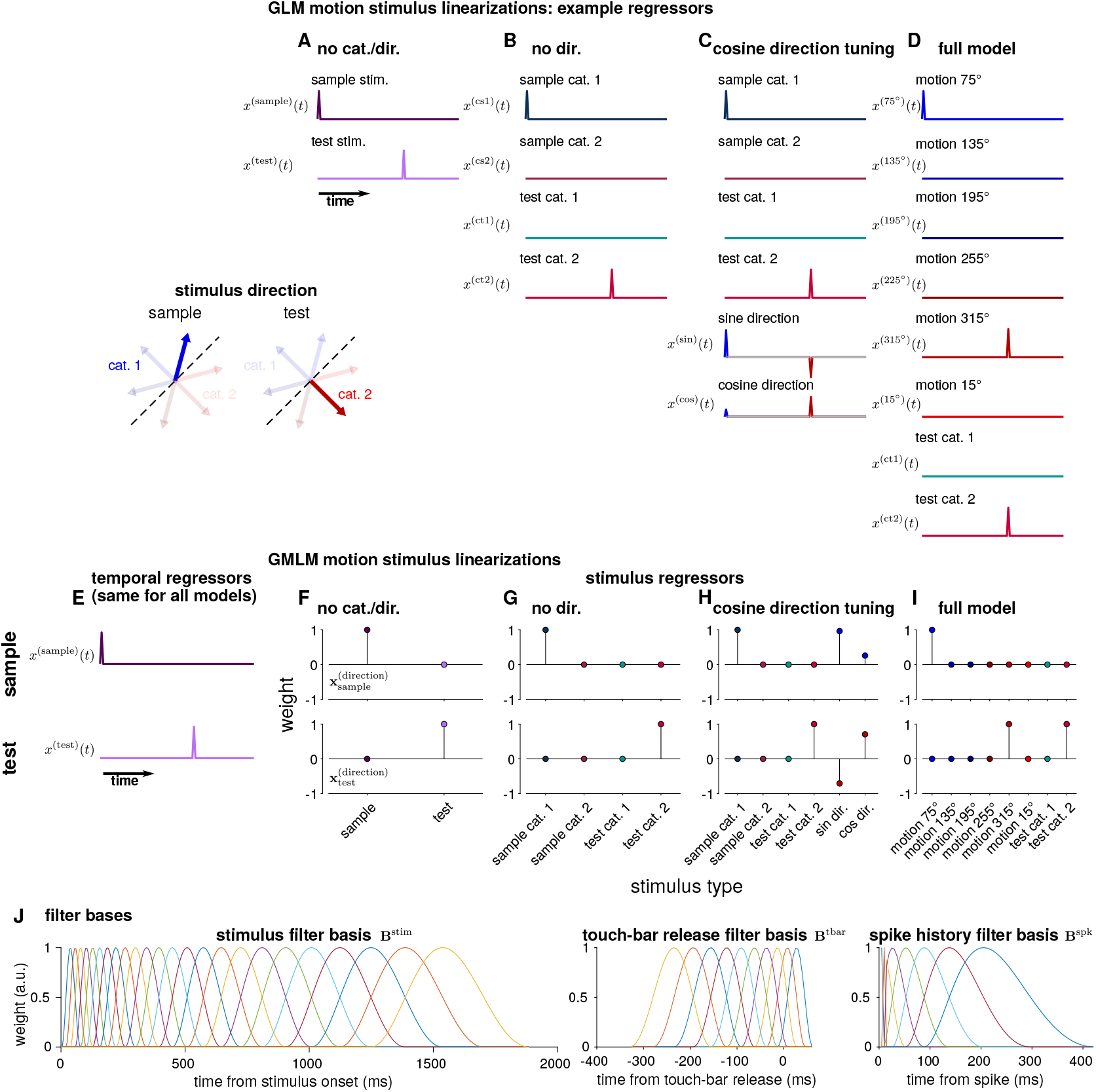
Linearizations of the DMS or DMC tasks in the GMLM. (**A-D**) The temporal event regressors for the four GLM types for an example trial with a sample stimulus 75° (category one) and test stimulus of 315° (category two). (**A**) The two stimulus events for the no category or direction tuning model. The top event is 1 at the sample stimulus onset time and 0 elsewhere, and the bottom event is 1 at the test stimulus onset time and 0 elsewhere. (**B**) The stimulus events for the no direction tuning model. The two sample (or test) category events encode the onset time of a sample (or test) stimulus only for a specific category (the sample category two event is 0 for this trial because the sample stimulus is category one). (**C**) The stimulus events for the cosine tuning model. The category events are the same as the category events in B. The sine (or cosine) event is equal to the sine (or cosine) of the stimulus direction at the onset of either stimulus. (**D**) The stimulus events for the full tuning model. The first six events are 1 at the onset time of a specific stimulus direction (sample or test). The two category events are the same as before. This configuration is identifiable while allowing the category tuning to be different between the sample and test period, while keeping the direction tuning constant. (**E-F**) The event regressors for the four GMLM types for the same trial configuration. (**E**) The temporal events for the sample (top) and test (bottom) stimulus onset times. (**F-I**) The GMLM stimulus weightings for the four model configurations for the sample (top) and test (bottom) stimulus correspond to the weight of the stimulus events in A-D. The complete temporal kernels in the corresponding GLMs are thus the outer product of the temporal regressors in E and the weights of the weights in F-I, summed over

**Figure S2:**
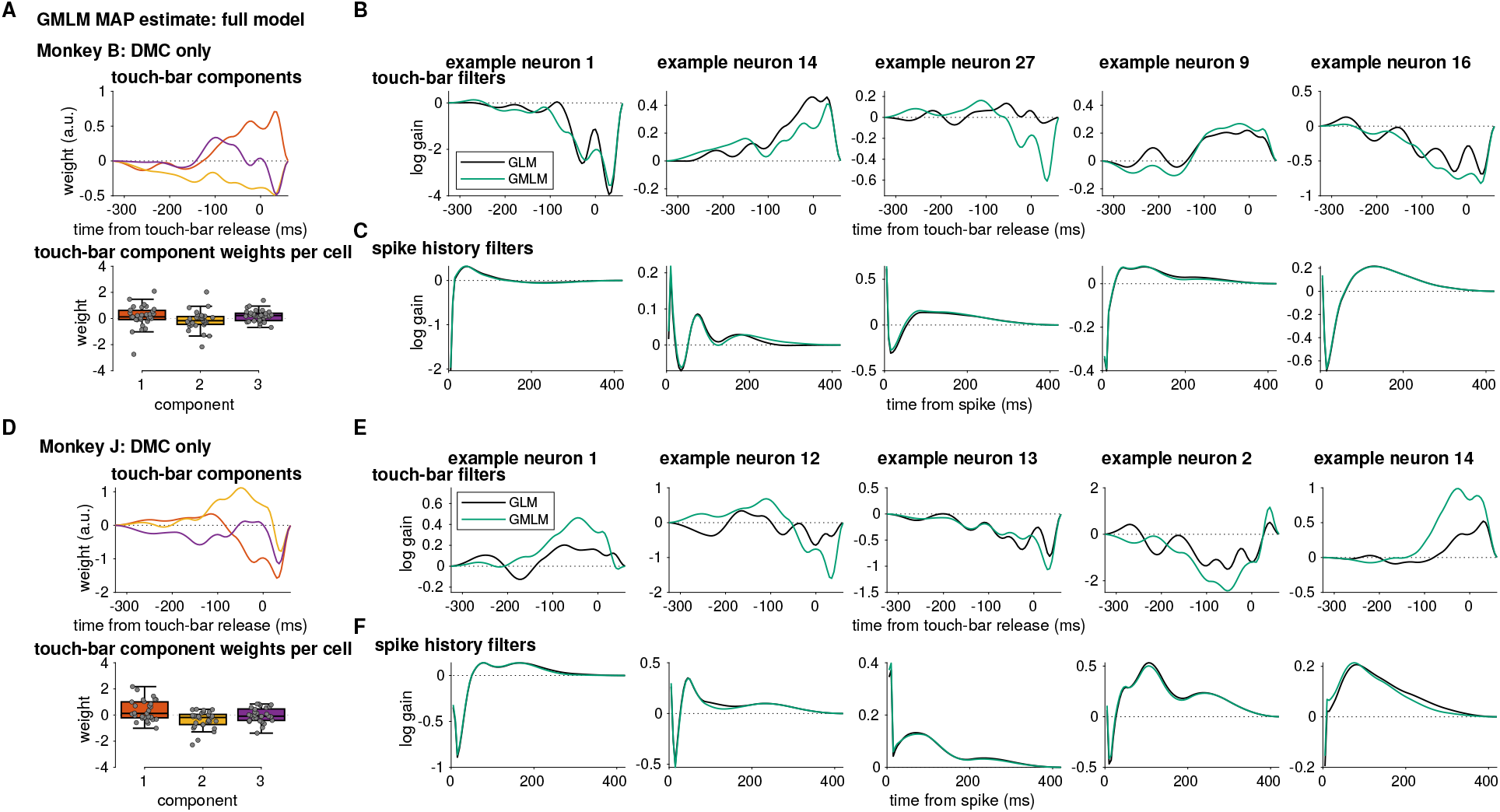
Touch-bar and spike-history kernels from the GMLM (full model) and the GLM fits. (A) The low-dimensional touch-bar release components for monkey B. (top) The three temporal kernels. (bottom) The loading weights for each touch-bar release component for each cell (points). Example touch-bar filters for five cells. The GLM touch-bar filters (black) are compared to the GMLM fit (cyan). (**C**) Spike-history filters fit to the same cells in B. The GLM spike-history filters (black) are nearly identical to the to the GMLM fit (cyan). (**D-F**) Same as A-C for monkey J.

**Figure S3:**
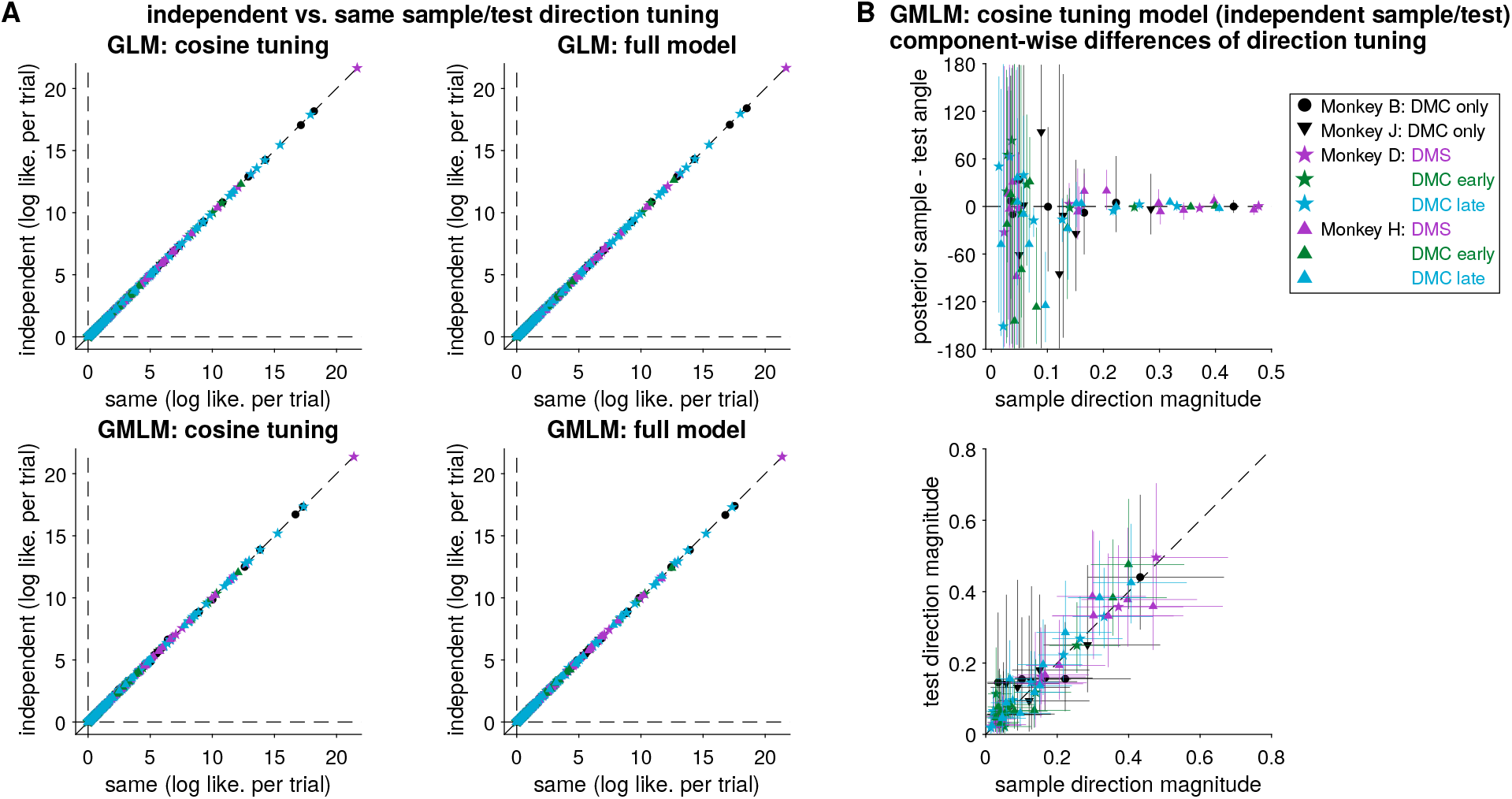
The GMLM and GLM find similar direction tuning across the sample and test stimuli. (**A**) cross-validated per-trial log likelihoods for each cell (relative to the GMLM without any stimulus terms, *R_s_* = 0). The left column shows comparisons using the cosine tuning model and the right column shows the full model. The top row compares the GLM fits to single cells and the bottom row shows the GMLM fits for the same model configurations. No population showed a significant improvement including the independent directions (for each model and population *p* > 0.8, one-sided Wilcoxin signed rank test with Benjamini-Hochberg correction). Several populations indicated that overfitting occurred with the GLM with independent directions: the same direction model was on average better. (**B**) Bayesian analysis of the cosine-tuned GMLM with independent sample and test direction parameters. Each point represents a single GMLM stimulus component for one population (i.e., there are seven points for monkey B because we selected seven GMLM stimulus components). (top) The posterior difference in preferred angle between the sample and test stimuli as a function of the magnitude of sample direction tuning. The angle is *θ*^(*r*)^ and the magnitude is *a*^(*r*)^ in Eq. 28 (see Methods). As the magnitude increases, the test and sample directions tend towards zero. At lower magnitudes, the preferred angle is difficult to estimate (undetectable) and therefore the difference shows high uncertainty. (bottom) The magnitude of the sample and test direction tuning for each component. Error bars show 99 % credible intervals.

**Figure S4:**
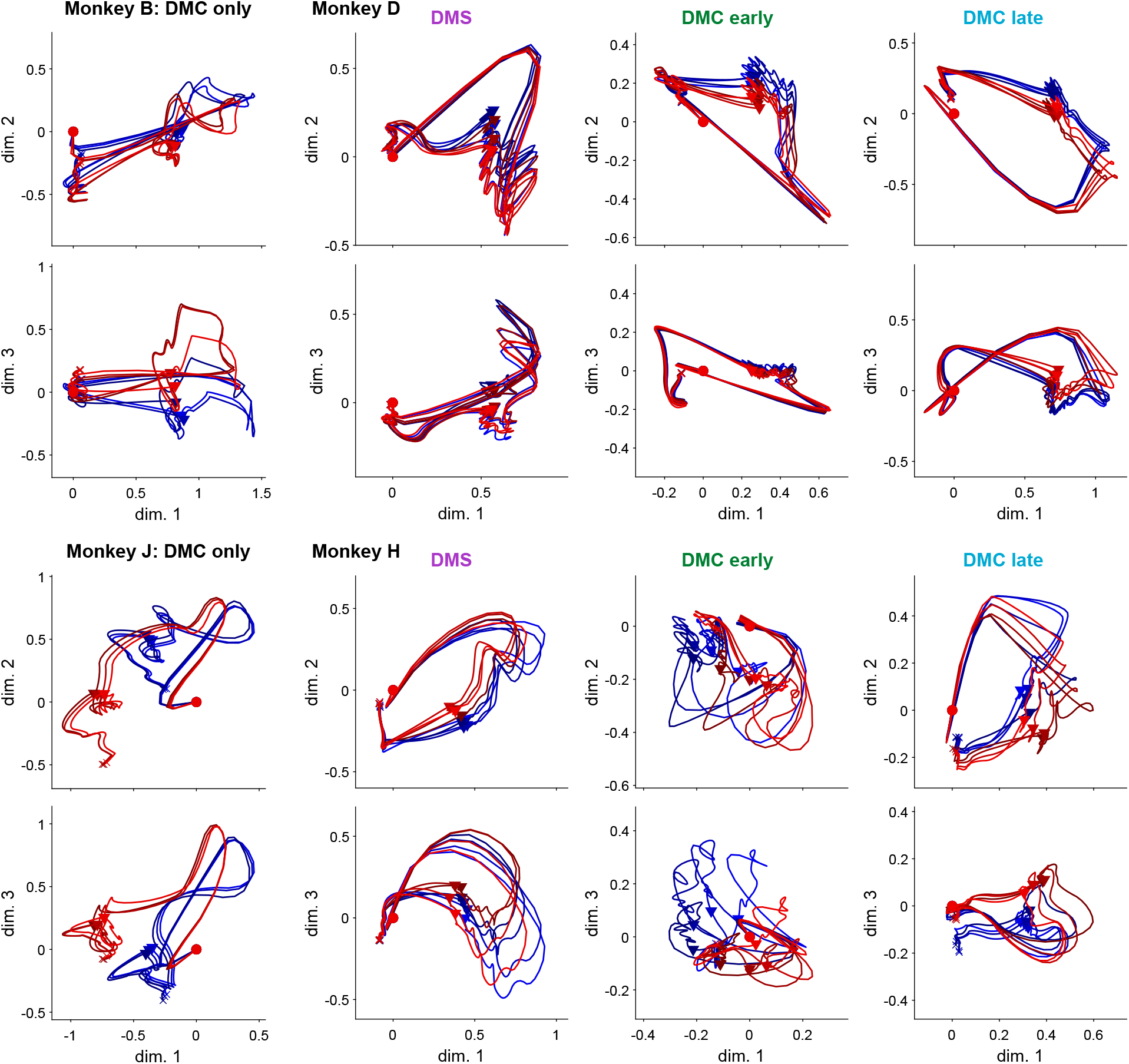
The top three dimensions of the GMLM subspaces (full model) in response to the sample stimulus for each animal and recording epoch without removing the mean over motions directions. Fig. 5 shows the subspaces and trajectories after removing the mean.

**Figure S5:**
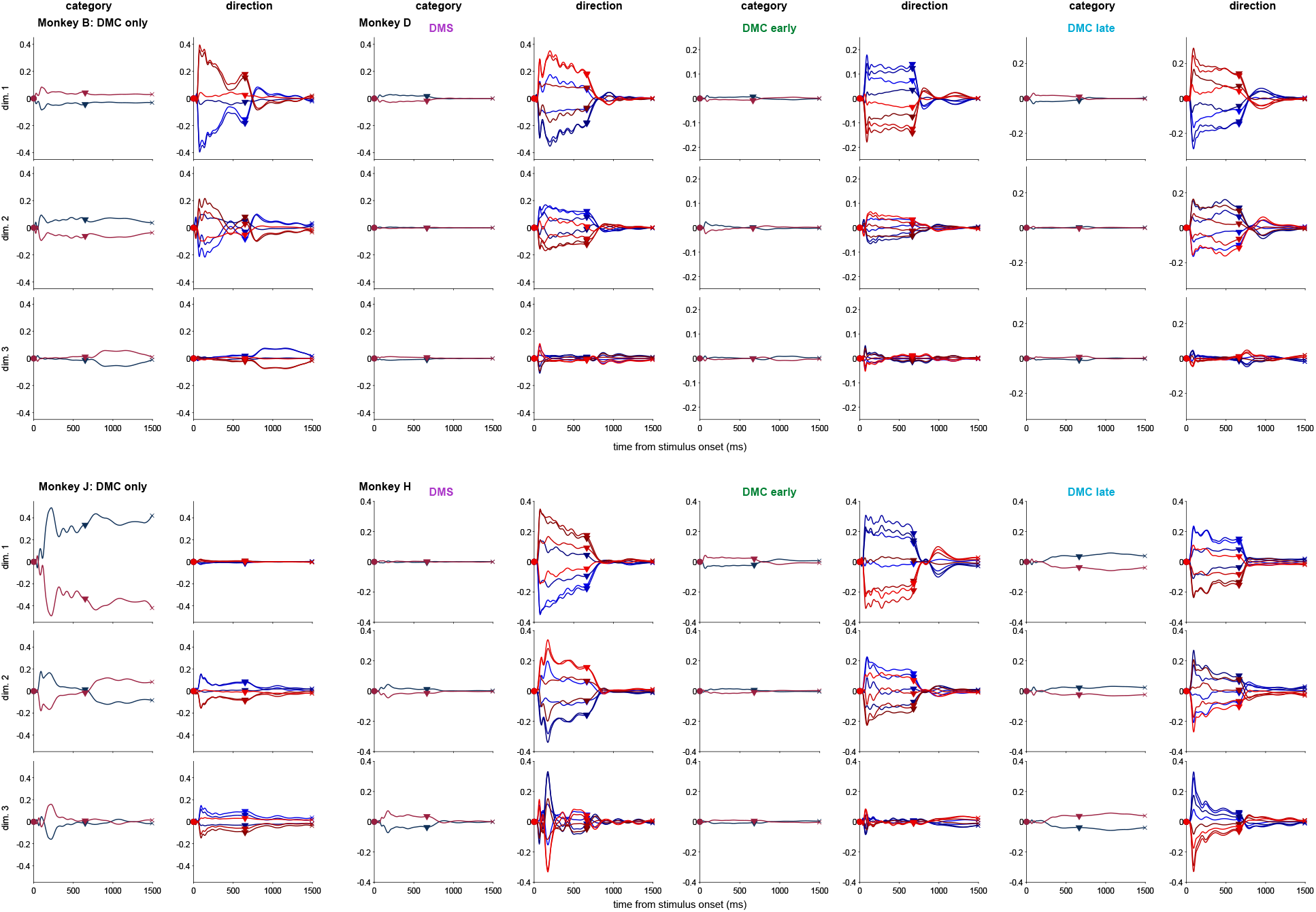
Low-dimensional subspaces of the GMLM with cosine direction tuning with the mean removed for each LIP population. Each of the top three dimensions are shown as a function of time relative to sample stimulus onset (this is similar to Fig. 5, but with the dimensions plotted separately relative to time). For the cosine model, we can separate the direction and category components. The left column for each population shows the sample category trajectories in the three dimensions. The right columns shows the direction trajectories, decoupled from category.

**Figure S6:**
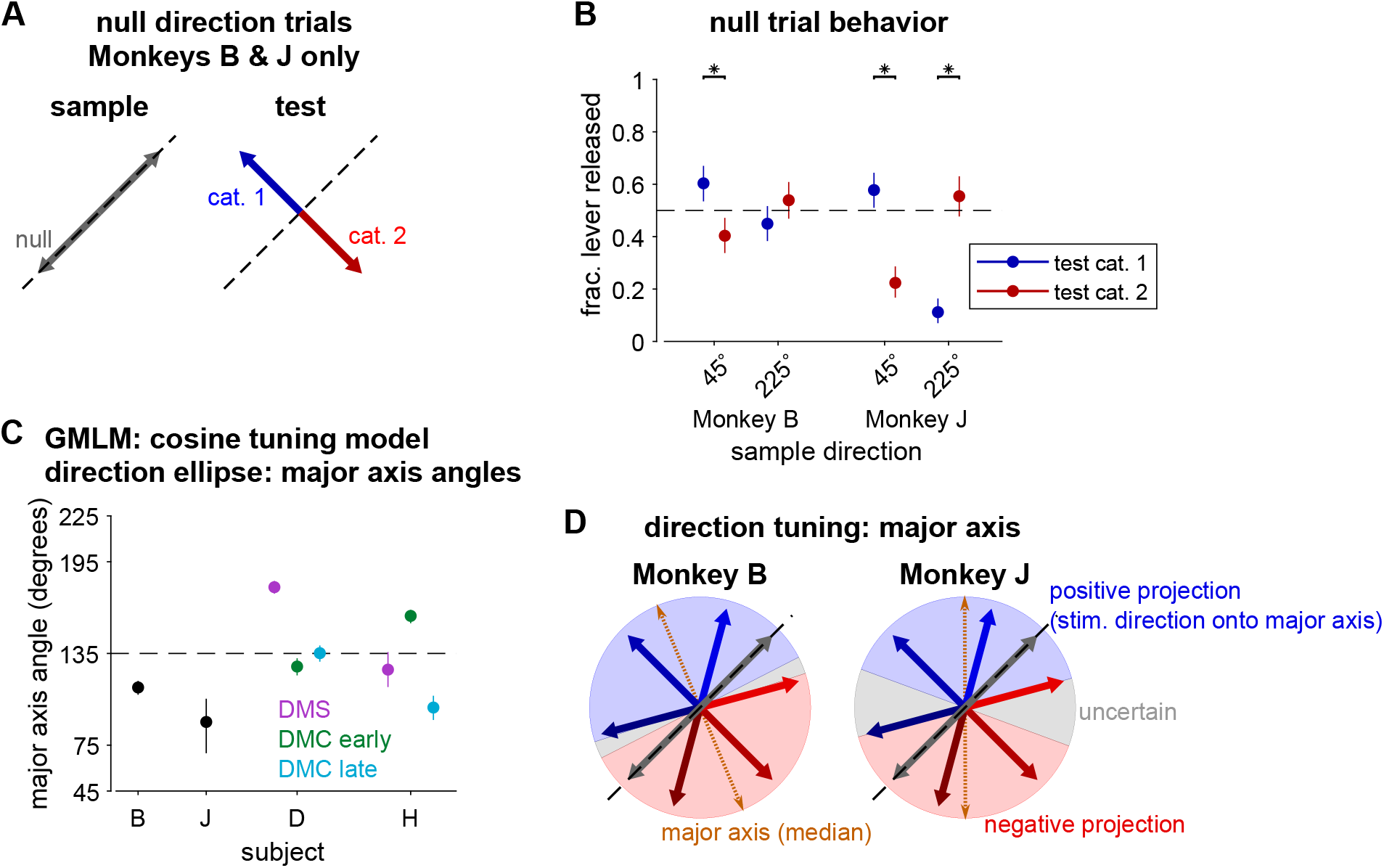
Analysis of the direction tuning of the low-dimensional GMLM cosine tuning model components. (**A**) The sample and test directions selected on “null” direction trials for monkeys B and J. The sample directions lie on the category boundary. These trials were not included in the GMLM analysis and were rewarded randomly. (**B**) Behavior for the four possible combinations of angles on the null-direction trials. The color indicates the test direction/category. The point shows the posterior mean estimate of the fraction of touch-bar released during the test stimulus presentation and the error bars denote a 99 % credible interval. Asterisks indicate that the response proportion for the two test directions was different for a given null sample direction (*p* < 0.01; two-sided Wilcoxon rank sum test, Holm-Bonferroni corrected). (**C**) Bayesian analysis of the cosine tuned GMLM. The GMLM defines the direction tuning as an ellipse in a low-dimensional space. We computed the angle of the major axis of the ellipse: the angle with the most modulation in the low-dimensional space (see Methods Eq. 26). The angle is only identifiable up to 180°. Therefore, we placed it within 45° to 225° to align with the task. If the axis aligned exactly with the categorization task, the angle would be 135°. The ellipse depends on time relative to stimulus onset, and so we took the mean angle during the first 650 ms of stimulus presentation. The points show the posterior median and the error bars denote a 99 % credible interval. (**D**) Illustration of how the direction ellipse’s major axis aligns with the task directions. The blue region shows where motion-direction angles project positively along the major axis vector (generally overlapping with category one). The red region shows where motion-direction angles project negatively along the major axis vector. The gray region shows angles that are within the 99 % credible region of the posterior (from C) and cannot be classified. We note that the regions do not exactly align with the category bounds. However, they do correlate with the monkeys’ choice biases for the null directions: for monkey B, the 45° null-direction (up and to the right) is in the blue region and the monkey was more likely to release the touch-bar on 45°trials when the test stimulus was in category one than for a category two test stimulus.

**Figure S7:**
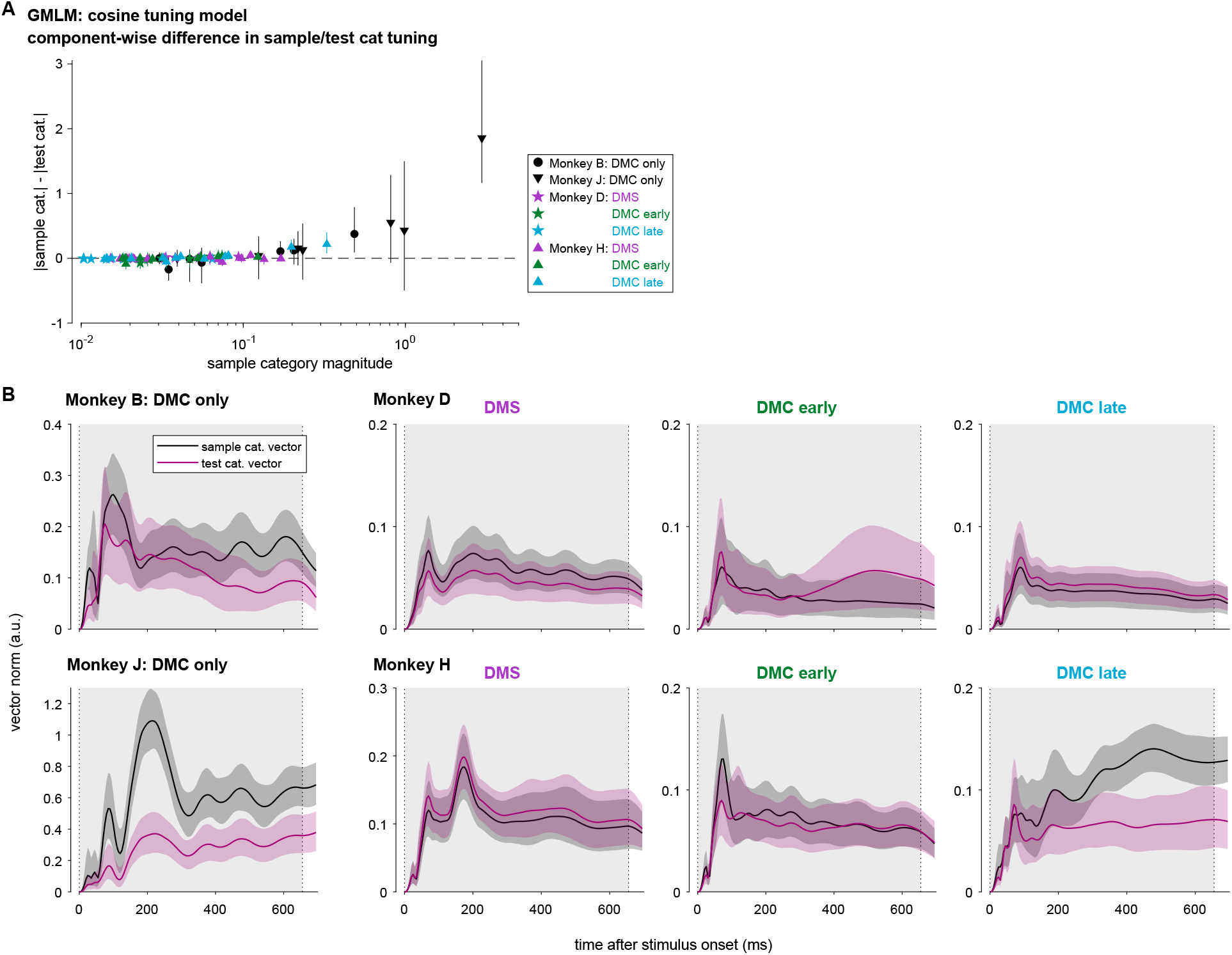
Analysis of the sample and test category tuning of the low-dimensional GMLM cosine tuning model components. (**A**) The difference in the magnitude of coefficients for the sample and test categories in the individual GMLM components (one point per each GMLM stimulus component per population). The component-wise sample category magnitude is computed as 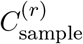 and the difference is 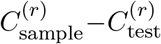 in Eq. 29 (see Methods). The points show the posterior median and the error bars denote a 99 % credible interval. (**B**) The norm of the category tuning vector as a function of stimulus onset time for the sample and test stimuli in each of the eight LIP populations. The category vector norm is given in Eq. 27 (see Methods). The traces show the posterior median and the shaded regions denote a pointwise 99 % credible interval.

**Figure S8:**
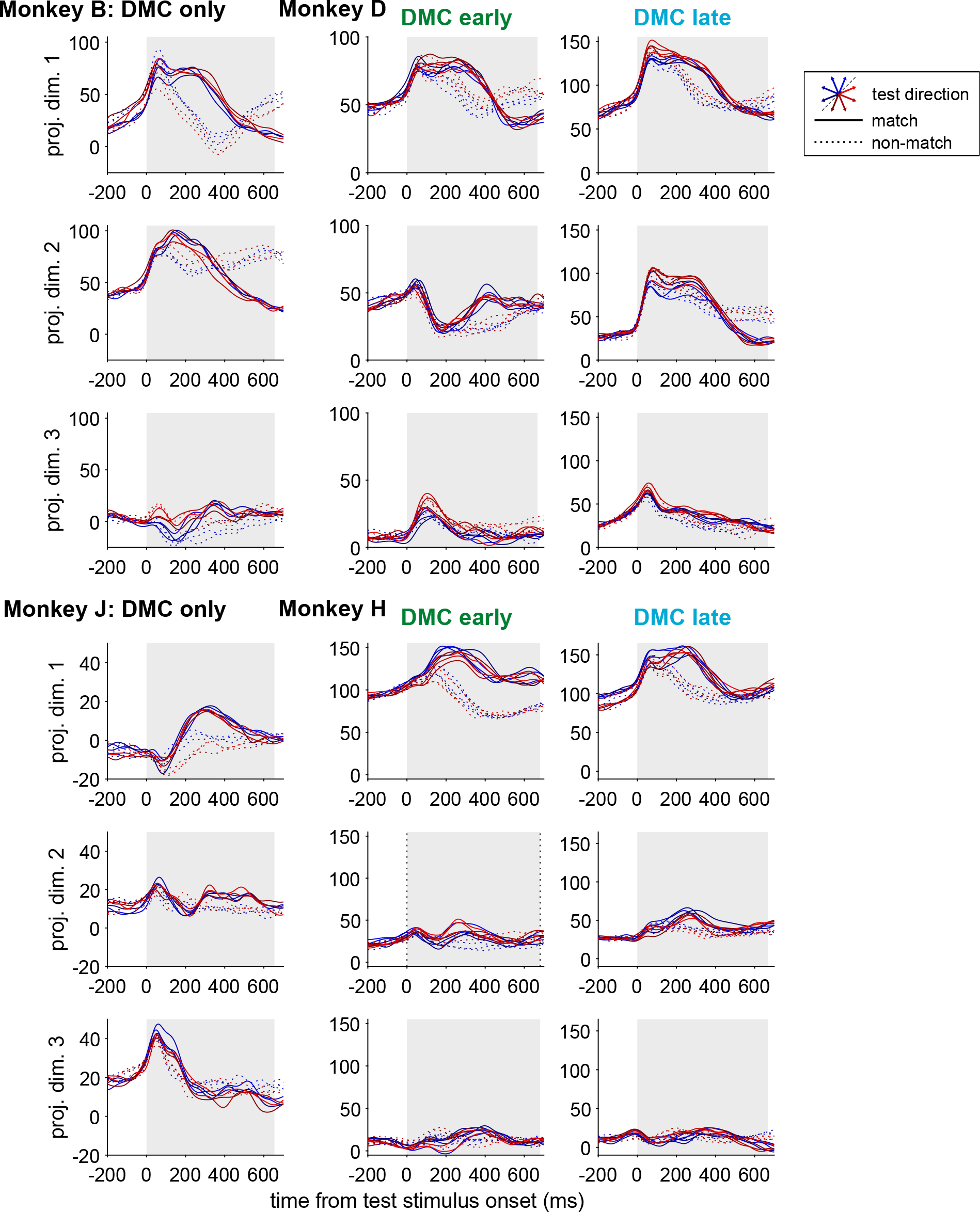
The PSTHs of the six DMC populations projected onto the three-dimensional touch-bar subspace fit by the full GMLM (the subspace given by orth(**V**^tbar^), see Methos). The PSTHs are conditioned by both test stimulus direction (color) and by match (solid lines) or non-match (dotted lines) trials. The gray region denotes the stimulus presentation period (although it is terminated early on match trials by the touch-bar release).

**Figure S9:**
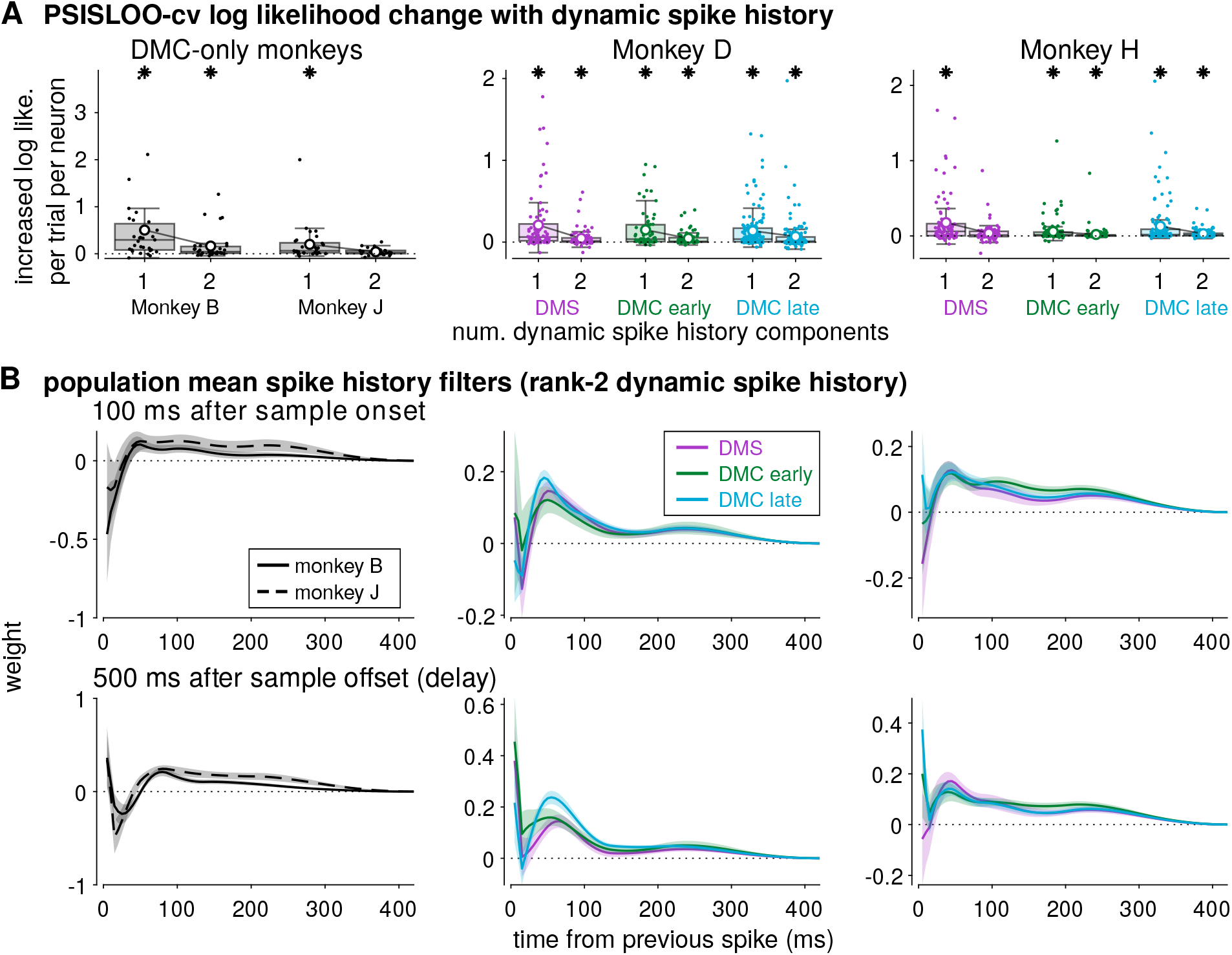
Including a dynamic spike history filter tensor improves model fit. (**A**) The mean change in cross-validated log likelihood per-trial for each neuron as a function of the number of components (i.e., the average improvement in predictive performance for adding an additional dynamic spike history component). The log likelihood for the rank-1 dynamic spike history is relative to the GMLM without any dynamic spike history (but still includes each individual neuron’s static spike history filters). Leave-one-out cross-validation was estimated for each trial using Pareto-smoothed importance sampling (PSISLOO-cv). Stars denote a statistically significant improvement after including the dynamic spike history component (*p* < 10^−4^, paired, one-sided Wilcoxin signed-rank test). The lines with white circle markers denote the average log likelihood per trial across neurons. While the improvement after including two dynamic spike history components was often statistically significant, it was less dramatic than the gain from a single component. (**B**) The mean population mean effective spike history filters for all eight LIP populations during sample stimulus presentation (top; 100 ms after stimulus onset) and during the delay period (bottom; 500 ms after stimulus offset). The spike history filters were computed as the MAP estimate of the GMLM with rank-2 dynamic spike history (*R*_*h*_ = 2). Error regions denote ±2 SEM.

